# Constitutive Interferon Epsilon Expression Shapes Antiviral Epithelial States in the Female Reproductive Tract and Intestine

**DOI:** 10.1101/2024.11.15.623843

**Authors:** Rebecca L. Casazza, Samantha Skavicus, David Hare, Kaila A. Cooley, Nicholas S. Heaton, Carolyn B. Coyne

**Author notes:** Address correspondence: Carolyn Coyne, PhD, 3130 Medical Sciences Research Building III, 3 Genome Court, Durham, NC 27710 USA.

## Abstract

Antiviral defenses at mucosal barriers are essential for preventing viral entry and systemic infection. Interferon epsilon (IFNε) is a unique type I IFN that, unlike other family members, is not induced by infection but is constitutively expressed in epithelial tissues. IFNε was initially characterized in the female reproductive tract (FRT), where it provides broad antiviral protection, but its roles outside the FRT remain poorly defined. Here, we used *Ifnε^-/-^* mice and single-cell RNA sequencing to delineate IFNε function across distinct mucosal surfaces. In the FRT, *Ifnε* expression was restricted to specific epithelial subsets, was independent of estrous stage, and maintained basal ISG expression. IFNε was also retained intracellularly in primary human FRT-derived cells. Extending these analyses to the intestine, we found that IFNε is highly expressed in villous-tip enterocytes of the small intestine *in vivo*, where it sustains inflammatory enterocyte subsets and maintains type III IFN expression. Loss of *Ifnε* depleted these subsets and rendered mice more susceptible to enteric viral infection. Together, these findings establish IFNε as a constitutively expressed, spatially restricted IFN that coordinates mucosal antiviral defenses across both reproductive and gastrointestinal epithelial tissues.

## Introduction

Antiviral defenses at barrier sites are essential for preventing entry and dissemination of viruses, which could compromise tissue integrity and lead to systemic infection. In the reproductive tract, these defenses are particularly crucial given exposure to external pathogens, such as sexually transmitted viruses, and their essential role in safeguarding reproductive health and ensuring successful reproduction. Similarly, in the intestine, antiviral programs are required to control constant exposure to enteric viruses while maintaining epithelial integrity and nutrient absorption. In both tissues, robust antiviral responses are needed to preserve a balanced immune environment, minimizing the risk of infection while sustaining barrier function and overall tissue homeostasis.

Interferon epsilon (IFNε) is a type I IFN that has been described as specifically enriched in the epithelial cells of the female reproductive tract (FRT) (Bourke et al., 2022; Fung et al., 2013). Unlike other type I IFNs, IFNε is constitutively expressed and is not induced by viral infections. Despite its lack of induction by infection, IFNε plays a direct role in innate antimicrobial defense against sexually transmitted infections, including HIV, herpes simplex virus (HSV), Zika virus (ZIKV), and Chlamydia (Coldbeck-Shackley et al., 2023; Fung et al., 2013; Stifter et al., 2018). Beyond antiviral defense, IFNε exerts tumor-suppressive activity in fallopian tube epithelial cells (Marks et al., 2023) and has been implicated in pregnancy, as it is expressed in the myometrium, cervix, and chorioamniotic membranes (Miller et al., 2022). Elevated levels of IFNε in amniotic fluid during spontaneous preterm labor with intra-amniotic infection suggest a role in antimicrobial defense within the amniotic cavity. However, consistent with viral infections, IFNε expression is not enhanced by bacterial stimulation *in vitro*, indicating that alternative cellular sources may contribute to its presence during infection(Fung et al., 2013; Miller et al., 2022).

Several clues exist as to the regulation of *IFNε* expression. Analysis of the *IFNε* promoter region revealed progesterone response elements implicating hormonal modulation in its expression (Hardy et al., 2004). Consistent with this, *Ifnε* expression fluctuates in the mouse uterus with the estrous cycle, and peaks during estrous when progesterone levels are highest (Fung et al., 2013). The transcription factor ELF3, which directly regulates the expression of genes important for maintaining the integrity and function of epithelial barriers, contributes to the control of IFNε expression in the uterus and levels fluctuate with estrous status (Fung et al., 2024). Transcription factor binding site analysis has also revealed potential regulatory sites for NF-κB, a critical mediator in immune and inflammatory responses (Hardy et al., 2004). NF-κB binding elements in the IFNε promoter suggest that this cytokine might be responsive not only to hormonal signals but also to cellular stress and inflammatory stimuli, which aligns with its role in epithelial immune defense. Indeed, treatment of cells with tumor necrosis factor-α (TNF-α) upregulates *IFNε* expression (Matsumiya et al., 2007). Taken together, these findings suggest that *IFNε* expression may be finely tuned by both hormonal and inflammatory cues, with transcription factors including NF-κB and ELF3 potentially coordinating epithelial immune readiness in response to estrous phase and/or local environmental stimuli.

In addition to antimicrobial and antitumor properties in the epithelium of the FRT, IFNε potentially plays a role in modulating the immune system. For example, *Ifnε^-/-^* mice exhibit less uterine NK cell accumulation at baseline and less activation after infection than WT mice, a phenotype which results in part from differences in cytokine secretion in *Ifnε^-/-^*macrophages/monocytes (Mayall et al., 2024). Additionally, T-, B-, and NK cells respond to recombinant IFNε *in vitro* (Stifter et al., 2018). These findings suggest that IFNε not only fortifies epithelial defenses but also plays a broader immunoregulatory role in the FRT, coordinating both innate and adaptive immune responses to maintain tissue homeostasis and readiness against infection.

Recent studies have suggested that IFNε might also play a role in inflammatory signaling outside of the FRT. In addition to epithelial cells of the FRT, *Ifnε* is expressed in the intestinal epithelium, where it helps limit inflammation and supports the intestinal regulatory T cell compartment (de Geus et al., 2024). In the testes, Ifnε is expressed in the germinal epithelium in spermatogenic cell, as well as in leydig cells and macrophages, where it protects against infection (Wijayarathna et al., 2024). Ifnε is also constitutively expressed in human primary bronchial epithelial cells, and in the intestines of bats (Deng et al., 2024; Kellner et al., 2025; Martinez-Espinoza et al., 2024). In rhesus macaques, IFNε is expressed in the epithelium of the large and small intestines, lung, and foreskin in addition to the female reproductive tract (Demers et al., 2014). These studies highlight IFNε as a multifaceted cytokine with roles in maintaining immune balance across diverse mucosal tissues, suggesting it may provide a widespread defense mechanism beyond the FRT.

The mechanisms regulating IFNε and its contributions to antiviral defense beyond the female reproductive tract (FRT) remain incompletely defined. To address this, we generated an IFNε knockout (*Ifnε*^-/-^) mouse using a modified intracytoplasmic gene editing via oviductal nucleic acids delivery (iGONAD) approach, which enables in vivo genetic modification without embryo transfer (Ohtsuka et al., 2018; Skavicus and Heaton, 2023). Consistent with a previously described *Ifnε*^-/-^ model (Fung et al., 2013), we confirmed that IFNε protects against intravaginal herpes simplex virus-2 (HSV-2) infection. Analysis of single-cell RNA sequencing (scRNA-Seq) datasets across the estrous cycle in the mouse FRT (Winkler et al., 2024) revealed that *Ifnε* is expressed in tissue-specific epithelial cell clusters and is not directly regulated by the estrous cycle. Instead, expression varies with fluctuations in the abundance of Ifnε-expressing epithelial subsets. These populations exhibited high interferon-stimulated gene (ISG) expression, which was absent in epithelial subsets lacking *Ifnε*, suggesting an autocrine mechanism of action. Supporting this, scRNA-seq of uterine tissues from WT and *Ifn^ε^*^-/-^ mice showed significant reductions in ISG expression specifically within Ifnε-expressing epithelial subsets, without broader transcriptional effects. In primary human FRT epithelial cells, IFNε was constitutively expressed, non-inducible by pathogens, and retained intracellularly. Similarly, IFNε was constitutively expressed and retained intracellularly in human stem cell-derived enteroids. Single-cell RNA sequencing of human enteroids and murine small intestine identified IFNε expression specifically in mature villous-tip enterocytes, establishing that IFNε is constitutively expressed within this epithelial subset *in vivo*. Mice lacking *Ifnε* exhibited reduced expression of antimicrobial genes, loss of inflammatory enterocyte populations that normally produce cytokines such as Ifnλ3, and increased susceptibility to oral infection with the enterovirus coxsackievirus B (CVB). These findings establish IFNε as a key contributor to mucosal immunity, sustaining antiviral defenses within tissue-specific epithelial cells of both the FRT and intestine, and broaden our understanding of its role beyond traditional pathogen-induced interferon responses.

## Results

### Generation of *Ifnε^-/-^* mice using iGONAD

We used a modified iGONAD protocol (Skavicus and Heaton, 2023) to generate an *Ifnε^-/-^* knockout mouse. The iGONAD technique was chosen for its efficiency and precision in gene editing, particularly in the context of creating knockout models. To achieve this, we employed a dual sgRNA approach targeting the single coding exon of the *Ifnε* gene, which resulted in a 263 basepair deletion (**Figure 1A).** Fifteen total pups were born following the injection and electroporation of sgRNAs and Cas9 into the oviducts of pregnant dams. These pups were subsequently screened for *Ifnε* knockout by PCR using primers spanning both gRNA target sites. The genotyping results revealed that 6 out of the 15 pups were homozygous for the *Ifnε* deletion, indicating a successful knockout of both alleles. Additionally, 3 pups were identified as heterozygous, carrying one wildtype and one knockout allele, while the remaining 6 pups were wildtype (representative genotypes, **Figure 1B**). Successful gene knockout was further validated by Sanger sequencing, which confirmed the deletion at the nucleotide level (**Figure Supplemental 1A**). Following this, a qPCR-based genotyping assay was developed in collaboration with a commercial vendor (Transnetyx) to streamline the genotyping process for subsequent generations. To establish a stable knockout line one animal (15) identified as homozygous for the *Ifnε* deletion was selected as the founding sire. To ensure genetic integrity and minimize potential off-target effects, this animal was backcrossed ∼6 times with C57Bl/6J mice. Following backcrossing, a stable and reproducible *Ifnε*^-/-^ line was developed and used for subsequent studies.

**Figure 1:**
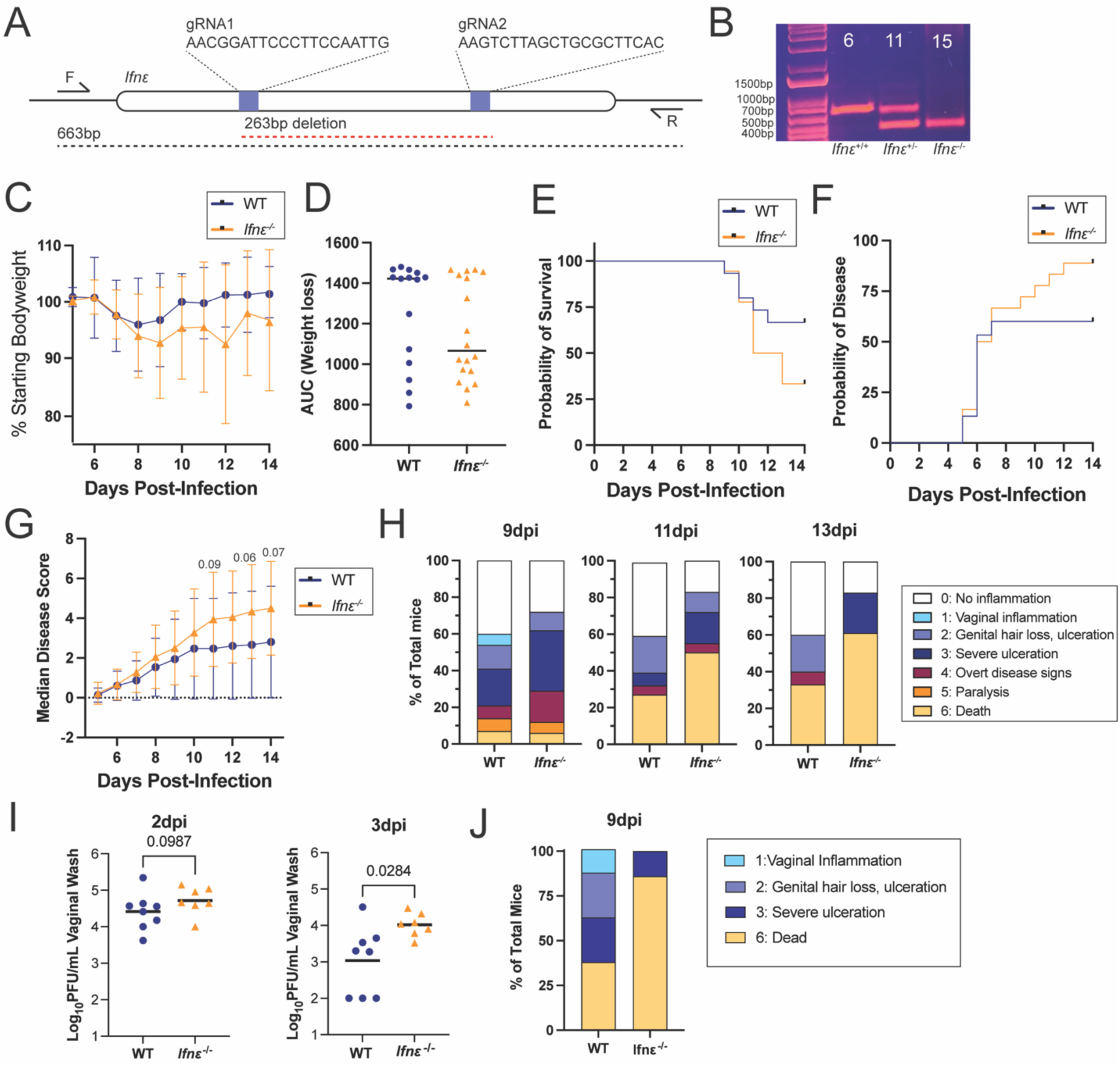
Increased HSV-2 Susceptibility in *Ifnε* Deficient Mice Generated via iGONAD CRISPR/Cas9. **A)** Schematic of CRISPR/Cas9 knockout strategy. **B)** Representative PCR genotyping results from the first generation of pups born to dams that underwent iGONAD gene editing. Mouse 6 (*Ifnε^+/+^),* 11 (*Ifnε^+/-^)* and 15 (*Ifnε^-/-^)* are shown. Mouse 15 was chosen as the founding sire for the line. **C-H)** WT (n=15) and *Ifnε^-/-^* (n=18) females were infected with 100PFU of HSV-2 333 and weighed and monitored for disease signs daily. Weight loss (C,D), survival (E), and disease onset (F) are shown. Disease onset (F) includes any observable pathology score. Median clinical score is shown in G, and in (H), frequencies of clinical scores are shown at multiple days post-infection (dpi) from 9-13dpi. Key at right. Differences in median clinical score was determined by Mann-Whitney test in Graphpad Prism. **I-J).** WT (n=8) and *Ifnε^-/-^* (n=7) females were infected with 10,000PFU of HSV-2 333. Vaginal lavages were collected at 2 and 3dpi and titrated by plaque assay. Significant differences were calculated by T-test in Graphpad Prism. Individual animals are depicted by symbols in (I). Frequencies of clinical scores are shown at 9dpi in (J).

To confirm that this newly established *Ifnε^-/-^* line recapitulated phenotypes observed in a previous *in vivo* model (Fung et al., 2013) regarding the role of IFNε in antiviral signaling in the FRT, we infected *Ifnε^-/-^* and WT female mice intravaginally with 100PFU of HSV-2 (strain 333) and scored mice daily for clinical signs. *Ifnε^-/-^* mice lost more weight compared to their WT counterparts (**Figure 1C, 1D**), a higher percentage of *Ifnε^-/-^* mice succumbed to infection (73% vs 40%), (**Figure 1E**) a higher percentage of *Ifnε^-/-^* mice had disease onset compared to WT mice (89% vs 60%), defined as any observable clinical sign, (**Figure 1F**) *Ifnε^-/-^* mice displayed higher frequencies of advanced clinical signs throughout the course of infection (**Figure 1G, 1H**). Despite *Ifnε^-/-^* having worse disease outcomes by every metric measured, none of these differences were significant, in part due to a large proportion of mice that remained uninfected. To increase infection rates in our mice, we performed infections with 10,000PFU of HSV-2 (**Figure 1I, 1J).** *Ifnε^-/-^* had higher viral burdens in vaginal lavages at 2 and 3dpi (**Figure 1I**) and had worse clinical outcomes measured by clinical scores (median of 3 in WT mice vs median of 6 in *Ifnε^-/-^* mice, p = 0.0741 measured by Mann Whitney) (**Figure 1J**). Taken together, these data show that *Ifnε^-/-^* mice generated using iGONAD recapitulate the heightened susceptibility and impaired antiviral response observed in a previous *in vivo* model of HSV-2 infection in the FRT.

### IFNε is a Key Modulator of Basal ISG Expression in the FRT Epithelium

To determine the specific cell types expressing *Ifnε* in the uterus and the functional consequences of *Ifnε* signaling, we performed scRNASeq on uterine tissue isolated from WT and *Ifnε^-/-^* mice in estrous, with 19227 cells passing quality control. Cluster analysis and cell-type specific marker expression following integration to correct for batch effects across independent animals revealed four distinct clusters of EpCs, two populations of endothelial cells (Endo), stroma and fibroblasts, two populations of muscle cells (Musc-1 and Musc-2), pericytes (Peri), and various immune cells including macrophages (Mac), dendritic cells (DC), and NK cells (NK) (**Figure 2A**). There were near-equivalent ratios of all cell types between WT and *Ifnε^-/-^*mice (**Figure 2B, Supplemental Figure 2A**) and cell clusters expressed cell type-specific markers, which were conserved between WT and *Ifnε^-/-^* mice (**Figure 2C**, **Supplemental Figure 2B**). We identified a single cluster of EpCs (EpC-2) that highly expressed *Ifnε* and that also expressed high basal ISG levels (**Figure 4D**). This cluster also expressed markers associated with LEpCs (e.g., *Prap1*, *Tacstd2*, *Ltf*) (**Supplemental Figure 2C**, **2D**).

**Figure 2.**
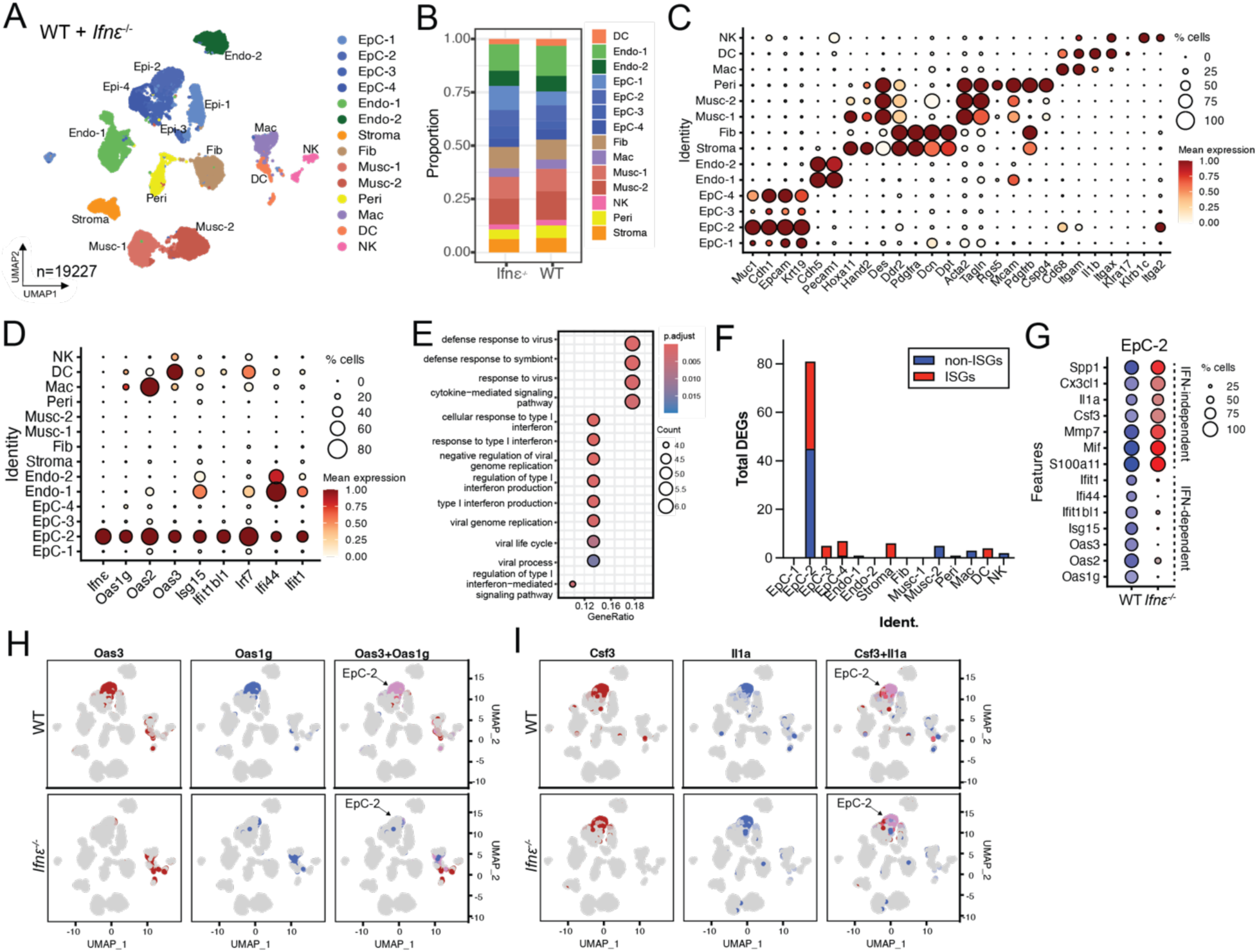
*Ifn*ε^−/−^ Mice Lack Basal ISG Expression in Uterine Epithelial Cells. **A)** UMAPs showing uterine cell clusters from *Ifn*ε*-/-* and WT mice. **B)** Bar plot of the proportions of each cell cluster in *Ifnε-/-* and WT mice. **C)** DotPlot of canonical cell type markers in each cell cluster. Key and scale at right. **D)** DotPlot of *Ifnε* and ISG expression in each cell cluster. Key and scale at right. **E)** Pathway analysis on differentially expressed genes in *Ifnε-/-* and WT uteruses. Shown are the gene counts per pathway and the p-value of each pathway. **F)** The total number of DEGS in each cluster. ISGs are shown in red and non-ISGs are shown in blue. **G)** DotPlot showing ISG and cytokine expression levels split by mouse genotype. Scale at right. **H)** FeaturePlots showing co-expression of *Oas3* and *Oas1g* in either WT (top) or *Ifnε^-/-^* (bottom) samples. **I)** FeaturePlots showing co-expression of *Csf3* and *Il1a* in either WT or *Ifnε^-/-^*samples.

**Figure 3:**
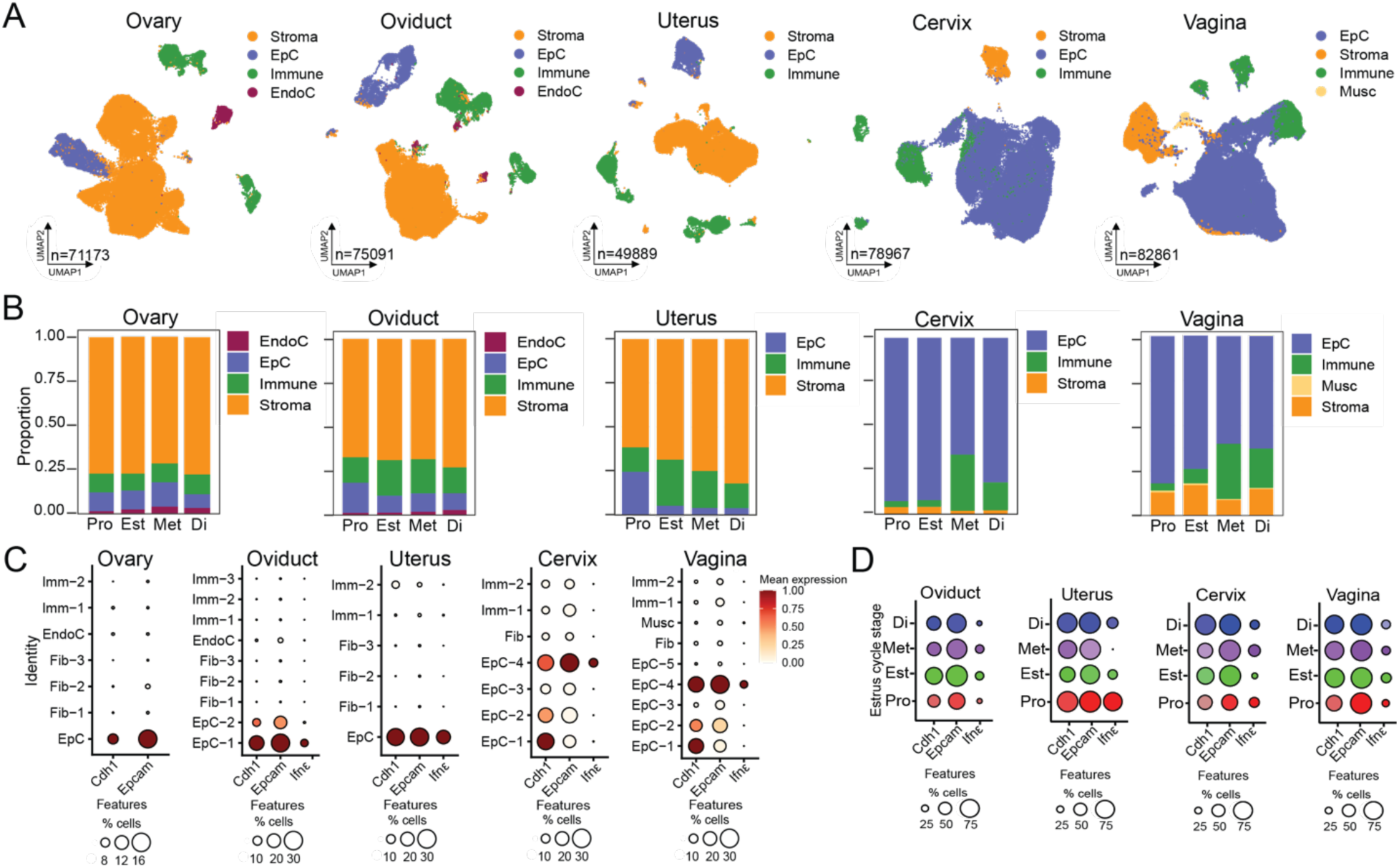
*Ifnε* is expressed in epithelial cells of the female reproductive tract throughout the estrous cycle. **A)** UMAPS showing clustering of stromal (Stroma, orange), epithelial (EpC, blue), immune (Immune, green), endothelial (EndoC, red), and muscular (Musc, yellow) cells within the different tissues of the female reproductive tract (FRT). **B)** The proportions of each cell type split by estrous stage: Proestrous (Pro); Estrous (Est); Metestrous (Met), and Diestrous (Di). **C-D)** DotPlots showing expression of epithelial cell markers and *Ifnε* in all clusters (C) or only in clusters that express *Ifnε* split by estrous stage (D).

**Figure 4:**
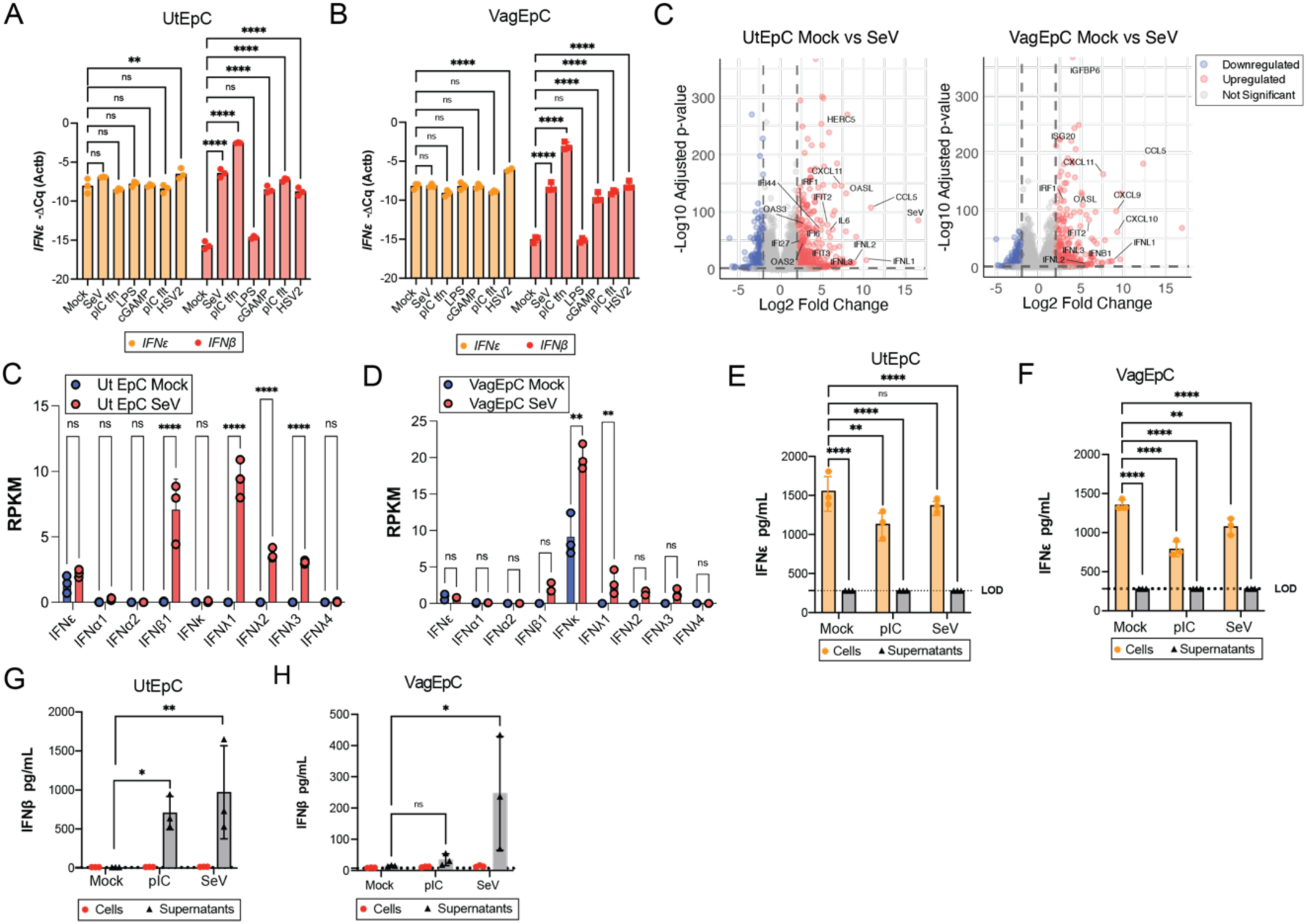
IFNε is retained intracellularly in primary human epithelial cells and is not induced by viral infections. **A, B),** Primary human uterine (UtEpV) (A) or vaginal (VagEpC) (B) epithelial cells were treated infected with Sendai virus cantell (SeV, 200HAU) or HSV2 (MOI of 1), treated with LPS or poly(I:C), or transfected with cGAMP or poly(I:C). Cells were harvested 20hpi and *IFNE* and *IFNB* levels were quantified by qRT-PCR. **C),** Volcano plots of differentially expressed genes (log2 ζ 2 or :: -2, padj ::0.01) in UtEpC (left) or VagEpC (right) infected for ∼24hrs with 200HAU SeV. Upregulated genes are shown in red and downregulated genes are shown in blue. Select gene names are shown in black. **D, E),** Reads per kilobase of transcript, per million mapped reads (RPKM) of the indicated IFN gene in Mock- (blue) or SeV-infected (red) UtEpC (D) or VagEpC (E). **F-I),** IFNε(F, G) and IFN-ý (H, I) protein levels were measured from cell lysates and cell supernatants from uterine (F, H) or vaginal (G, I) epithelial cells by ELISA. Significance was determined by two-way ANOVA using a Tukey’s test for multiple comparisons (*, p<0.05, **p<0.01, ***p<0.001, ****p<0.0001, ns, not significant).

To assess whether *Ifnε* expression was required for basal ISG expression in EpC-2 and other cell types, we performed pseudobulk differential expression analysis across all cell clusters in WT and *Ifnε*^-/-^ mice. We identified a total of 115 differentially downregulated transcripts in *Ifnε^-/-^* mice compared to WT, with ∼70% of these differentially expressed genes (DEGs) attributed specifically to the EpC-2 cluster (**Supplemental Figure 2E**). Pathway analysis of these DEGs showed a significant enrichment in genes associated with IFN signaling and the antiviral response (**Figure 2E**). Of the 81 differentially downregulated DEGs in EpC-2, ∼45% of them were ISGs (**Figure 2F).** Several additional clusters also exhibited a downregulation of select ISGs (EpC-3, EpC-4, Stroma, and DC), but the total number of DEGs was significantly lower than that observed in EpC-2 (**Figure 2F**). Additional clusters, including EpC-1, Endo-2, Fib, and Musc-1 did not exhibit any changes in gene expression in *Ifnε^-/-^* mice (**Figure 2F**). There were only two DEGs enriched in *Ifnε^-/-^* mice, *Ppbp* and *Crct.* Importantly, although ISGs were significantly downregulated in EpC-2, other non-IFN-mediated inflammatory transcripts were unchanged, including those associated with IL1 signaling (e.g., *Il1a*), chemokine signaling (e.g., *Cx3cl1*), and cytokine signaling (e.g., *Mif*) (**Figure 2G-2I**). Taken together, these data show that *Ifnε* expression is crucial for maintaining basal ISG expression in EpC populations in the FRT.

### *Ifnε* is expressed in epithelial cells of the oviduct, uterus, cervix, and vagina independent of estrous status

To determine to determine the impact of the estrous cycle on the expression of *IFNε* in the FRT, we analyzed an existing single cell atlas of the cycling mouse female reproductive tract based on 378,516 cells from normal cycling young mice in the four cycle phases (proestrus [Pro], estrus [Est], metestrus [Met], and diestrus [Di]) (Winkler et al., 2024). The dataset contained cells harvested from ovary, oviduct, uterus, cervix, and vagina from 3-5 independent animals. We restricted our analyses to non-pregnant young mice (aged <12 months). A total of 71173 (Ovary), 75091 (Oviduct), 49889 (Uterus), 78967 (Cervix), and 82861 (Vagina) cells passed quality control and were used for cluster analysis and cell-type specific marker expression analysis following integration to correct for batch effects (**Figure 3A, Figure S3A-J**). We broadly clustered cells into epithelial cells (EpC), stromal cells, immune cells, endothelial cells (EndoC), or muscle cells. Each tissue site exhibited site-specific enrichment of these cell types, consistent with what has been described (Winkler et al., 2024), with the cervix and vagina most enriched in EpCs and ovary, oviduct, and uterus more enriched in stromal cells (**Figure 3A**, **Figure S3A-J**). While there was slight variation in the proportion of these cell types during the estrous cycle, including the enrichment of immune cell types in the cervix and vagina during Met, the proportion of cell types were largely maintained across all tissues (**Figure 3B**).

We next determined the cell type(s) expressing *Ifnε* across all tissue sites and defined the impact of the estrous cycle on this expression. For this analysis, we further clustered cell types to increase resolution. In the oviduct, uterus, cervix, and vagina, we found that *Ifnε* was specifically enriched in EpC cell types that also expressed high levels of epithelial markers (e.g., *Cdh1*, *EpCam*) (**Figure 3C**). We did not detect any *Ifnε* in the ovaries (**Figure 3C**). We next determined the levels of *Ifnε* expressed in EpCs across the estrous cycle in all tissues. In the oviduct, cervix, and vagina, we observed highly similar levels of expression across all stages of the estrous cycle (**Figure 3D**). In the uterus, levels of *Ifnε* were consistent across Pro, Est, and Di, with significantly lower levels in Met (**Figure 3D**). Similarly, *Ifnε* levels were low in the uterus of pregnant mice (**Figure S3K, S3L**). Taken together, these data indicate that IFNε is predominantly expressed in epithelial cell types across the FRT, with consistent expression levels across the estrous cycle, except in the uterus where levels significantly decrease during Met and pregnancy.

### *Ifnε*-expressing epithelial cells in the FRT express high basal ISGs

The FRT is lined by specialized epithelial cell types, each with distinct functions tailored to their anatomical location. These include ciliated and secretory EpCs in the oviduct, luminal and glandular EpCs in the uterus, mucus-secreting EpCs in the cervix, and stratified squamous EpCs in the vagina. Given that *Ifnε* was expressed in EpC populations across the FRT, we next sought to identify the specific EpC populations responsible for this expression and whether it varied by location. Furthermore, we sought to investigate whether *Ifnε*-expressing epithelial cell populations exhibited distinct transcriptional profiles, including elevated ISG levels. To do this, we subsetted the EpC cell cluster from all anatomical sites and re-clustered to obtain better resolution between distinct EpC types. In the oviduct, EpC cells were clustered into ciliated (cEpC) and secretory (SEpC) populations, which were distinguished based upon marker expression (**Supplemental Figure 3A-3B**). We found that *Ifnε* was specifically expressed in an SEpC cell population marked by *Serpina1e* expression (**Supplemental Figure 3C, 3D**). This population also exhibited high basal levels of ISGs (**Supplemental Figure 3C**). In the uterus, EpCs were distinguished between markers associated with luminal EpCs (LEpCs) and glandular EpCs (GlEpCs) (**Supplemental Figure 3E**), with *Ifnε* expression restricted to a LEpC population expressing high levels of Tactstd2 (**Supplemental Figure 3F-3H**). This cluster also expressed high levels of diverse ISGs (**Supplemental Figure 3G**). The cervix contained the largest number of EpCs, which sub-clustered into intermediate (IEpCs), superficial (SuEpCs), and basal EpCs (BEpC), some of which were proliferating (pBEpC) (**Supplemental Figure 3I, 3J**). *Ifnε* was expressed in both SuEpC populations, with significantly higher levels in the Sprr2d^+^ cluster, which also showed elevated ISG expression (**Supplemental Figure 3K, 3L**). The vaginal EpCs clustered into SuEpCs, BEpCs, and columnar EpCs (ColEpCs), with ColEpCs expressing both *Ifnε* and basal ISGs (**Supplemental Figure 3M-3P**). These data demonstrate that *Ifnε* is expressed in specific epithelial cell populations across tissue sites, with its expression correlating with high basal levels of ISG expression within these clusters.

The data described above suggested that *Ifnε* is expressed in distinct EpC types in an anatomical location-specific manner, suggesting that these cell types might exhibit conserved expression of factors responsible for *Ifnε* expression. To determine if this was the case, we compared the differentially enriched genes in all *Ifnε* expressing clusters across anatomical sites and identified the genes shared between these cell types. This analysis revelated that only 31 genes (1% of the total genes) were shared between *Ifnε* expressing EpC types (**Supplemental Figure 5A**). The top-most gene shared across all cell clusters was *Elf3* (**Supplemental Figure 5B**), which has been shown previously to regulate *Ifnε* expression (Fung et al., 2024). In fact, there were only two transcription factors shared between all *Ifnε*-expressing EpC clusters, which included *Elf3* and *Maff* (**Supplemental Figure 5C**). These findings suggest that, despite the anatomical specificity of *Ifnε* expression, only a small set of conserved transcription factors, which included *Elf3*, may play a role in regulating *Ifnε* expression across diverse epithelial cell types.

### IFNε is retained intracellularly and is not induced by viral infections in primary human epithelial cells

To assess whether IFNε expression levels changed following infection, we cultured human primary endometrial and vaginal epithelial cells, infected them with Sendai virus (SeV) or HSV-2, or treated them with a variety of PAMPs including poly(I:C), cGAMP, and LPS, and measured *IFNε* expression by qRT-PCR. Consistent with what we observed in the mouse scRNAseq analysis described above, *IFNε* was basally expressed in both uterine and vaginal primary cells and expression levels were unchanged with infection or PAMP treatment (**Figure 4A, 4B)**. In contrast, *IFNβ* was undetected at baseline, and was induced by all conditions except for LPS treatment (**Figure 4A, 4B).** To broadly profile the induction of *IFNε* and other IFNs, we performed whole transcriptome bulk RNA-Seq on uterine and vaginal epithelial cells infected with SeV, as well as on mock-infected controls. Both cell types responded to infection by induction of various IFNs, including *IFNβ* and *IFN*λ 1-3, as well as canonical ISGs (e.g., *OAS1*, *OASL*, *ISG15*) (**Figure 4C-4E**). In contrast, although *IFNε* was expressed basally, it was not induced by SeV infection in either cell type (**Figure 4D, 4E**).

In our scRNAseq datasets, we observed basal ISG expression in clusters that also expressed *Ifnε*, suggesting an autocrine role for IFNε signaling in subsets of epithelial cells of the FRT. To determine if IFNε is retained intracellularly, we examined both lysates and supernatants from human uterine and vaginal primary epithelial cells for IFNε or IFNβ levels by ELISA. IFNε was undetectable in the supernatants from vaginal and uterine primary cells basally, after poly(I:C) treatment, and after SeV infection (**Figure 4F, 4G**). However, there were high levels of IFNε detected in the cell lysates in all conditions, although there was significantly more IFNε at baseline than there was following poly(I:C) treatment or infection (**Figure 4F, 4G).** In contrast, IFNý was undetectable at baseline and although it could be observed at low levels in cell lysates following poly(I:C) treatment or SeV infection, it was significantly higher in the supernatants **(Figure 4H, 4I).** Together, these data indicate that IFNε is constitutively expressed and retained intracellularly within epithelial cells of the FRT, where it likely functions through autocrine signaling, in contrast to IFNβ, which is secreted in response to infection.

### IFNε is basally expressed in and retained intracellularly in human intestinal organoids

IFNε has been reported to be expressed in the gastrointestinal tract and may be associated with colitis (de Geus et al., 2024). To determine if IFNε is retained intracellularly in the gastrointestinal tract as well as in the FRT, we examined cells and supernatants of human stem-cell derived enteroids. To include diverse GI cell types, human enteroids were grown in expansion media (Exp) to maintain a proliferative state with stem and progenitor cells or under differentiation conditions (Dif) to promote their maturation into specialized cells, including enterocytes, goblet cells, and Paneth cells (Fujii et al., 2018; He et al., 2022). Consistent with the FRT epithelium, IFNε was undetectable in the supernatants of both Exp and Dif enteroids and was only detected by ELISA in the cell lysates of both culture conditions (**Figure 5A**). To define the expression of IFNε in the GI tract we performed scRNASeq of intestinal organoids under differentiation and expansion conditions. Enteroids cultured under these conditions contained CD24^+^ progenitor cells, transit amplifying cells, immature enterocytes, mature enterocytes, BEST4^+^ ionocytes, goblet cells, and enteroendocrine cells, which expressed canonical markers associated with these cell types (**Figure 5B-5D**). As expected, we found that Exp enteroids were enriched in CD24^+^ cells, transit amplifying cells, and immature enterocytes and Dif enteroids were enriched in mature enterocytes, and BEST4^+^ ionocytes and exhibited differences in cell type-specific expression (**Figure 5C, Supplemental Figure 6A, 6B**). We found that *IFNε* was enriched in both mature enterocyte populations, as well as in a population of immature enterocytes (**Figure 5D, Supplemental Figure 6C**). The only other IFN detectable in the dataset was *IFNκ*, which was expressed at very low levels in CD24^+^ cells (**Figure 5D**).

**Figure 5:**
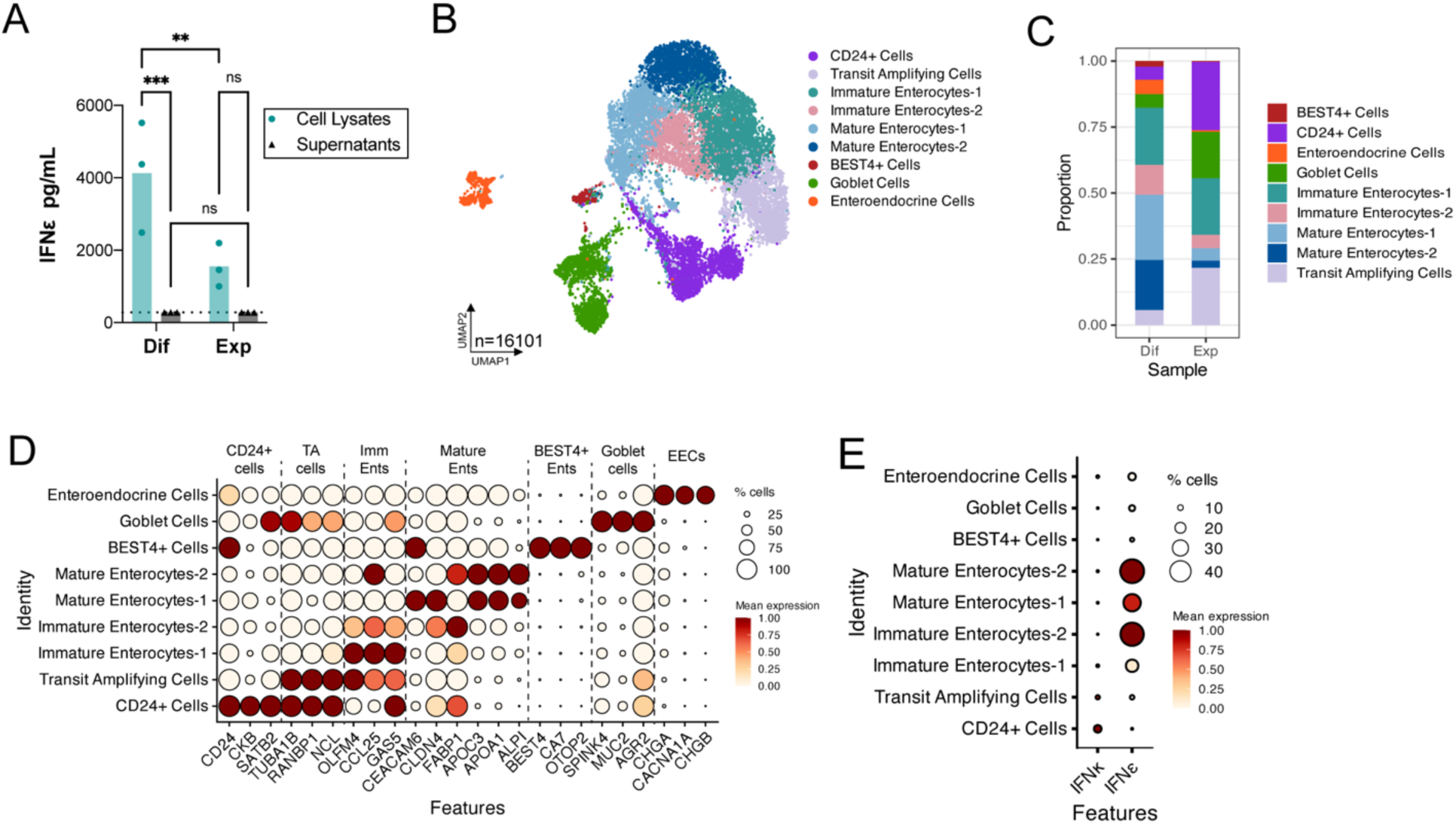
IFNε is expressed in human intestinal epithelial cells and is retained intracellularly. **(A-E),** Human enteroids were grown under expansion (Exp) or differentiating conditions (Dif) and then analyzed by scRNAseq. **A)** IFNε levels in supernatants and cell lysates from Exp and Dif enteroids, measured by ELISA. Data points represent IFNε concentration across experimental conditions, with significance determined by two-way ANOVA followed by multiple comparisons to assess differences between groups (ns, not significant, **p<0.01, ***p<0.001). **B)** UMAP plot depicting distinct cell clusters identified in the dataset. Each cluster is represented by a unique color, indicating cellular heterogeneity and grouping based on transcriptomic profiles. **C)** Bar plot depicting cell enrichment under Exp or Dif conditions. **D),** DotPlot of canonical cell makers used to define clusters. Key and scale at right. **E)** *IFNε* and *IFNκ* expression in cell clusters.

### IFNε is expressed in proximal, villus-tip enterocytes in the small intestine

To define the cellular expression patterns of *Ifnε* in the small intestine and assess its role in regulating ISG expression in the GI tract, we performed scRNA-seq on whole small intestines from WT and *Ifnε^-/-^* mice. All samples were merged and integrated together to correct for batch effects and to ensure identification of cell types across experimental conditions, with 93,309 cells passing quality control. Clustering identified 23 unique clusters (**Figure 6A).** All clusters were present across all samples and there were similar proportions of each cluster in WT and *Ifnε^-/-^* samples **(Supplemental Figure 7A, 6B).** Clusters were annotated based on the expression of conserved marker genes, identifying major epithelial lineages (enterocytes, Paneth cells, goblet cells, tuft cells, enteroendocrine cells, and stem cells), as well as immune (T and B cells) and mesodermal populations **(Figure 6B).** Consistent with our observations in enteroids, we observed that *Ifnε* expression *in vivo* was primarily restricted to enterocyte populations, with Enterocyte cluster 2 (Ent2) exhibiting the highest proportion of *Ifnε* positive cells **(Figure 6C, 6D).** Because only ∼10% of cells within Ent2 expressed *Ifnε*, we subsetted this cluster and performed reclustering to resolve enterocyte subpopulations and more precisely define the epithelial subset responsible for *Ifnε* expression (**Figure 6E).** Reclustering resolved 7 transcriptionally distinct enterocyte subpopulations, among which Ent2-D showed the highest *Ifnε* expression, with 60% of cells expressing the gene (**Figure 6F).** Ent2-D was marked by expression of duodenal cytochrome B (*Cybrd1*), a ferric reductase that is expressed in the brush border of duodenal enterocytes (McKie et al., 2001), as well as *Nt5e*, which is associated with villous tips (Moor et al., 2018) **(Figure 6G)**. These features indicate that *Ifnε* expression is enriched in proximal small intestinal enterocytes at the villous tips. To investigate this further, we calculated a villous index score for each cell type in the intestinal dataset, defined as the ratio of villous tip-associated genes to the sum of villous tip- and villous bottom-associated genes, as described previously (Moor et al., 2018). The *Ifnε*-expressing subset of Ent2 displayed relatively high villous index scores, indicating enrichment in enterocytes positioned near the villous tips **(Supplemental Figure 7C).** This localization is consistent with previous observations that *Ifnε* is preferentially expressed in epithelial cells situated closest to the luminal surface of the colon (de Geus et al., 2024). To assess whether *Ifnε*-expressing cells also exhibit elevated ISG levels, we performed a correlation analysis between *Ifnε* and all other genes within the subsetted Ent2 dataset (**Figure 6H**). No significant correlations were observed between *Ifnε* and ISGs, suggesting that *Ifnε* may serve functions in the small intestine distinct from basal antiviral gene induction. Instead, the top genes positively correlated with *Ifnε* included *Syt8*, *Cldn4*, *Add2*, and *Mucl3*, markers associated with villous tip enterocytes (**Figure 6H–I**).

**Figure 6:**
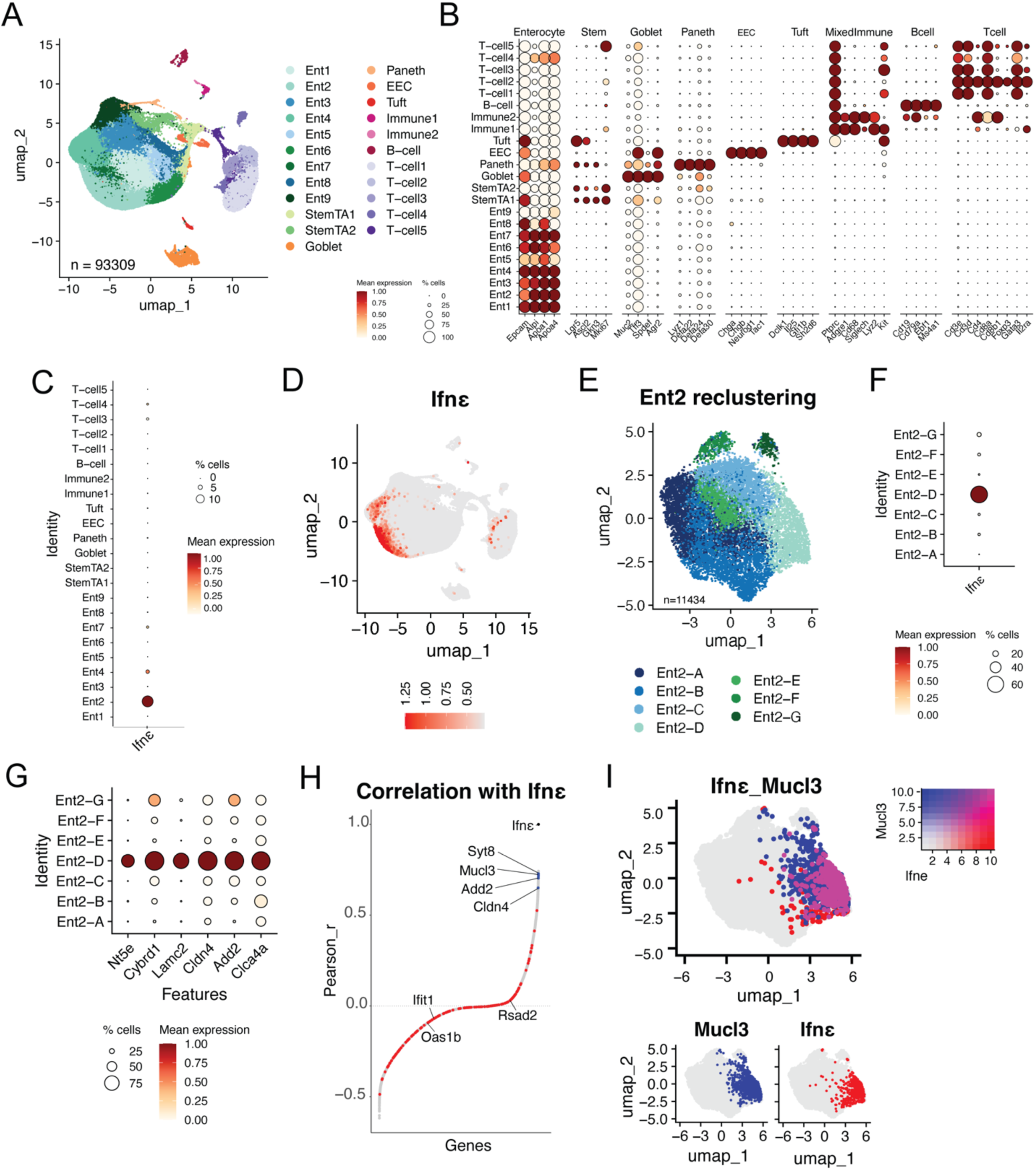
*Ifnε* is expressed in villous-tip enterocytes of the mouse small intestine. Single-cell suspensions were generated from WT and *Ifnε*^-/-^ mouse small intestines and analyzed by scRNA-seq. **(A)** UMAP plot revealed distinct cell clusters, including enterocytes (Ent), stem or transit-amplifying cells (Stem/TA), enteroendocrine cells (EEC), and immune cell populations expressing macrophage, dendritic cell, and mast cell markers. **(B)** Canonical marker genes used to define each cluster are shown in a dot plot. **(C, D)** *Ifnε* expression was detected primarily in enterocytes, as shown in both dot plots (C) and feature plots (D). **(E)** To further resolve enterocyte heterogeneity, cells within Ent2 were subsetted and reclustered, revealing distinct subclusters. **(F)** *Ifnε* expression across these subclusters is shown in a dot plot. (G) Dotplot showing expression of top markers in Ent2-D, the cluster that expresses *Ifnε* **(H)** Correlation analysis between *Ifnε* and all other genes expressed in Ent2 demonstrated no association with interferon-stimulated genes (ISGs, shown in red) but revealed strong correlations with villous tip–associated genes, including *Mucl3*. **(I)** A feature plot confirmed the overlap between *Ifnε* and *Mucl3* expression within Ent2, indicating that *Ifnε* is enriched in villous-tip enterocytes.

### Loss of *Ifnε* depletes inflammatory enterocyte subsets and increases susceptibility to enteric viral infection

To determine the broader impact of loss of *Ifnε* expression on intestinal homeostasis, we compared the small intestinal transcriptomes of WT and *Ifnε^-/-^* mice. Using pseudobulk differential expression analysis across all clusters in the dataset, we identified genes differentially expressed between genotypes (**Figure 7A).** DEGs were most abundant in enterocyte clusters, where they were enriched in pathways involved in antimicrobial and immune responses **(Figure 7B).** Notably, genes such as *Reg3g* (antimicrobial lectin), *Nos2* (nitric oxide synthase), *Tifa* (NF-κB signaling adaptor), and *Alpk1* (innate immune kinase) were differentially expressed across multiple enterocyte subsets **Figure 7C, Supplemental Figure 8A, 8B).** To further resolve enterocyte heterogeneity, we subsetted and reclustered Ent1-8, excluding Ent9, which was defined by mitochondrial gene expression and likely represented dying cells. This analysis revealed a distinct population of inflammatory enterocytes (NFκB^Ent^) present in WT intestines but largely absent from *Ifnε*^-/-^ samples (**Figure 7D–E**). The NFκB^Ent^ cluster was characterized by expression of *Nfkbia* (encoding an NFκB inhibitor), the NFκB regulator *Ubd*, and multiple inflammatory mediators, including *Nos2*, *Ccl20*, and *Cxcl11* (**Figure 7G**). Although NFκB^Ent^ cells did not express *Ifnε*, they were the only enterocyte subset to express *Ifnλ3*, the gene encoding a member of the type III IFN family (**Figure 7G**). *Ifnκ* was the only other IFN detected in enterocytes, but it was not expressed in NFκB^Ent^ cells. Strikingly, *Ifnλ3* expression was readily detected in WT enterocytes yet was completely absent in *Ifnε^-/-^* samples, suggesting that *Ifnε* may be involved in maintaining Ifnλ expression in the intestinal epithelium.

**Figure 7:**
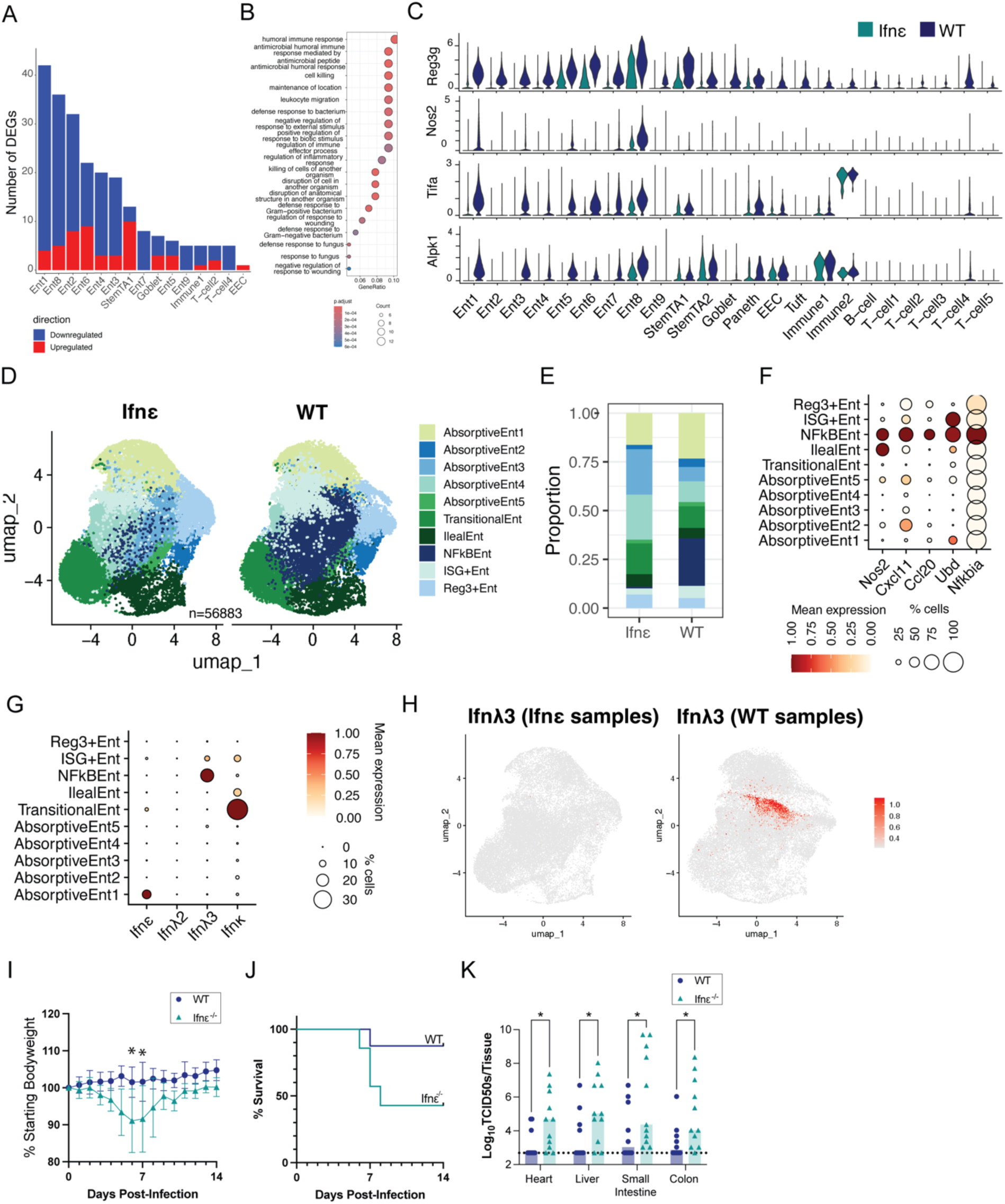
Loss of *Ifnε* Depletes Inflammatory Enterocyte Subsets and Increases Susceptibility to Enteric Viral Infection. **A)** Pseudo bulk analysis using DESEQ2 was performed for each cluster in the entire tissue intestinal single cell RNAseq data set introduced in Fig. 6. Total number of DEGs (in *Ifnε^-/-^* samples compared to WT) are shown for each cluster. Clusters not graphed (B-cell, Immune2, Paneth, STEMTA2, T-cell1, T-cell3, T-cell5, Tuft) had no DEGs. **B)** GO analysis was performed on the list of total identified DEGs and pathways identified were graphed. Scales and are below. **C)** Violin plots of gene expression of DEGs identified inmultiple clusters split by WT (dark blue) and *Ifnε^-/-^* (teal) samples. **(D-H)** All cells within enterocyte clusters 1-8 (Enterocyte cluster 9 was excluded as its top markers were all mitochondrial genes) were subsetted from the original dataset and reanalyzed. **D)** Split UMAP depicting the clustering of the Ent1-9 subcluster. **E)** Bar plot showing the proportion of cells in each cluster in either WT or *Ifnε^-/-^* samples. **F)** Dot plot showing the expression of top markers of the NFkB Enterocyte cluster. Key and scale below. **G)** Dot plot showing expression of all interferons detected in the Ent1-9 subcluster. **H)** Feature plot showing expression of *Ifnλ3* in either *Ifnε^-/-^* (left) or WT (right) samples. Scale to the right. **(I-K)** WT and *Ifnε^-/-^* mice were orally infected with 1x10^8^ PFU of CVB-H3 and weighed and monitored daily for signs of illness (I,J), or sacrificed at 4 days post infection (dpi). Viral burdens in tissues (K) were determined by TCID50. Significant differences in weight (*p<0.05) were calculated by t-test in Graphpad Prism. Differences in viral burden were calculated by Mann-Whitney test (*p<0.05).

As Ifnλ is a central mediator of antiviral protection in the GI tract (Lazear et al., 2019), we next tested whether *Ifnε*^-/-^ mice were more susceptible to enteric viral infection. To test this, we infected WT and *Ifnε^-/-^* mice with coxsackievirus B (CVB), an enterovirus that infects via the enteral route. Mice were infected with CVB3 (H3) by oral gavage. *Ifnε*^-/-^ mice exhibited significantly greater weight loss than WT controls (**Figure 7I**) and showed increased mortality following infection (**Figure 7J**). Notably, viral burdens in secondary organs, including the heart, liver, pancreas, and brain, sites commonly affected by CVB, were markedly elevated in *Ifn*ε^-/-^ mice relative to WT controls (**Figure 7K**). To confirm that these differences were not due to microbiome alterations in WT or *Ifnε^-/-^*mice, which have been shown to impact CVB infection (Robinson et al., 2019), we conducted 16S rRNA sequencing of the small ribosomal subunit from fecal samples of *Ifnε^+/-^* and *Ifnε^-/-^* littermates, thereby controlling for potential cage effects. The resulting dataset contained 2,695 OTUs, excluding those with a total count below three. On average, 76,503 quality-filtered reads were generated per sample, ensuring a robust analysis of microbiome composition (**Supplemental Figure 9A)**. High-quality reads were classified using the Silva v. 138 reference database, and OTUs were aggregated at each taxonomic rank (**Supplemental Figure 9B**). Alpha diversity, measured by the Shannon index, was comparable between *Ifnε^-/-^* (mean = 4.004, SD = 0.015) and WT (mean = 3.865, SD = 0.262) mice (**Supplemental Figure 9C)**. Differential abundance testing using DESeq2 revealed no OTUs with statistically significant differential abundance between WT and *Ifnε^-/-^* mice, indicating that microbiome composition was similar across genotypes. Together, these findings indicate that *Ifnε* protects against enteric viral infection, thereby acting as a critical barrier that prevents systemic viral dissemination from the gastrointestinal tract.

## Discussion

Antiviral defenses at mucosal surfaces are a crucial first-line defense against systemic infection. Type I IFNs play well-established roles in mucosal immunity and are induced by infection to elicit antiviral programs in target cells by signaling through the type I IFN receptor. Although IFNε is a type I IFN that also signals through the type I IFN receptor, it is not induced by infection or PAMPs. IFNε was initially characterized in the epithelium of the FRT where it was proposed to be hormonally regulated, and more recently in the epithelium of the colon, indicating that IFNε has a role in immunity across many epithelial barriers. Here, we demonstrate that IFNε plays a constitutive, essential role in antiviral defenses across mucosal sites. Using an *Ifnε^-/-^* mouse model, we confirmed that IFNε provides innate antiviral protection within the FRT, where it is expressed in specific epithelial cell clusters in a location-dependent manner and is associated with basal ISG expression independent of estrous cycle. Extending these findings to the intestine, we showed that IFNε is expressed in villous-tip enterocytes of the small intestine, where it supports inflammatory enterocyte subsets and sustains *Ifnλ3* expression. Loss of *Ifnε* led to depletion of these subsets, impaired *Ifnλ3* expression, and heightened susceptibility to enteric viral infection with systemic dissemination. Together, these results establish IFNε as a critical component of mucosal immunity, providing sustained antiviral defense across both reproductive and gastrointestinal epithelial tissues. Together, these results highlight IFNε as a critical component of mucosal immunity, offering sustained antiviral defense across both reproductive and GI epithelial tissues.

IFNε has been primarily studied in the context of the FRT, where it is expressed in the epithelium and has direct antimicrobial and antitumor activity. Using scRNAseq, we further defined the epithelial populations that express *Ifnε* throughout the mouse FRT using an existing single cell atlas of normally cycling mice (Winkler et al., 2024). We found that *Ifnε* is expressed in distinct epithelial cell types in each tissue: luminal and secretory cells in the uterus; columnar cells in the vagina; ciliated cells in the oviducts; and superficial cells in the cervix. In each epithelial population that expressed *Ifnε* we examined its expression levels at each estrous cycle stage. While *Ifnε* expression showed some fluctuations with estrous stage within individual tissues, it was consistently observed at all stages, with no patterns of fluctuation that were consistent across all tissues. The cell types that express *Ifnε* in the uterus; secretory and luminal epithelial cells, make up the layers of the endometrium that expand in preparation for embryo implantation during estrous (Jin, 2019) which could result in the higher gross *Ifnε* transcripts at a tissue level that others have observed in the uterus. These findings refine our understanding of *Ifnε* expression in the FRT, revealing its presence in distinct epithelial populations across tissues and estrous stages, suggesting a stable role in mucosal immunity that adapts to tissue-specific epithelial dynamics, especially in the endometrial layers preparing for implantation.

Previous studies have identified a role for IFNε in modulating inflammation in colitis models and shown Ifnε expression by IHC in the colon (de Geus et al., 2024). To further define the cell types that express IFNε in the intestine, we analyzed *IFNε* expression in human enteroids and in mouse intestinal scRNAseq data sets. We used enteroid models cultured under two separate conditions to capture the spread of intestinal cell types in the intestine and found *IFNε* in mature and immature enterocytes in enteroids. In mice we observed *Ifnε* primarily in enterocytes, as well as in select T-cell populations. Murine enterocytes enriched for *Ifnε* also expressed markers of villus-tip localization as well as a duodenal marker, indicating that *Ifnε* expression is restricted to proximal villous-tip enterocytes. These epithelial cells are the ones most directly exposed to luminal contents, positioning IFNε to provide front-line antiviral protection. Unlike in the FRT, where Ifnε expression correlated with ISG enrichment, intestinal analyses revealed no such association. Instead, *Ifnε*^-/-^ mice showed loss of key antimicrobial genes, including *Nos2* and *Reg3g*, across multiple enterocyte clusters. Strikingly, ∼25% of enterocytes in WT mice formed an immune-activated cluster characterized by expression of chemokines and *Ifnλ3* that was nearly absent in *Ifnε^-/-^* mice. This reveals a previously unrecognized crosstalk between constitutive IFNε and inducible type III IFNs, whereby IFNε sustains epithelial *Ifnλ* expression programs essential for gut antiviral immunity. Prior studies have shown that tonic IFNλ signaling establishes a basal antiviral state that restricts enteric pathogens (Van Winkle et al., 2022). Our findings suggest that IFNε functions as an upstream regulator of this tonic IFNλ environment, maintaining baseline antiviral readiness in the intestinal epithelium. In accordance with this, In *Ifnε^-/-^* mice we observed that IFNε exhibits antiviral activity against the enteric virus coxsackievirus B (CVB) following oral infection. *Ifnε^-/-^*mice exhibited increased weight loss and mortality following infection which corresponded with higher viral titers in the intestines, heart and liver at 4dpi. Taken together, these data define a role for IFNε throughout the intestines, where it is expressed in enterocytes and controls viral infection.

It is possible that IFNε is expressed at other cellular barriers including the upper and lower airway, blood brain barrier, urinary tract, testes and/or eyes. We initially expected that IFNε- expressing cell types would share common features; however, we observed distinct anatomical differences among the epithelial populations expressing *IFNε*. IFNε has been identified in tissues by IHC and is often found in the epithelial cells lining the lumen of tissues (de Geus et al., 2024; Demers et al., 2014; Fung et al., 2013; Marks et al., 2023). This pattern suggests that IFNε expression may be influenced by specific environmental or spatial triggers within these barriers. Recently, the transcription factor ELF3 was identified as driver of *Ifnε* expression in the mouse uterus (Fung et al., 2024), where it appears to support tissue-specific immune readiness. ELF3 is an epithelium-specific transcription factor involved in regulating the expression of genes essential for barrier function and epithelial integrity. While we consistently observed *Elf3* expression almost exclusively in *Ifnε*-expressing clusters throughout the FRT, in the intestine *Elf3* was detectable in nearly all cell types, indicating that there are other factors that control *IFNε* expression. This finding indicates that distinct transcriptional programs may regulate IFNε expression across different tissues, potentially adapting its expression to localized immune needs and environmental cues. Together, these observations suggest that IFNε expression is not only anatomically specialized but may also be regulated by diverse transcriptional drivers in response to the unique environmental contexts of each barrier tissue.

We focused on the direct antiviral effects of IFNε signaling in epithelial cells, but it is likely that IFNε also plays a nuanced role in immune cell signaling. Many groups have reported that immune cells respond to recombinant IFNε treatment by becoming activated, expanding, or upregulating ISGs (de Geus et al., 2024; Garcia-Minambres et al., 2017; Marks et al., 2019). This is not surprising, as type I IFNS can activate or influence virtually all immune cell types. It has been observed that IFNε knockout mice have reduced NK cells numbers in the uterus, a phenotype that results in part from decreases in IL-15 signaling in *Ifnε^-/-^* macrophages/monocytes (Mayall et al., 2024). In mouse models of colitis, *Ifnε*^-/-^ mice exhibit decreased levels of Tregs in the intestines which exacerbates intestinal pathology (de Geus et al., 2024; Fung et al., 2013). These findings indicate that IFNε likely has a role in regulating resident tissue immune cells as well as those that respond during infection. In our scRNAseq data set of the mouse intestine, we observed that a small portion (1-2%) of two our T-cell clusters expressed *Ifnε* and that several immune cell clusters, including T-cells, had differentially expressed genes in our *Ifnε^-/-^* mice that included cytokines such as *Il17a*. It remains to be determined if these differences in immune cells contribute to the phenotypes we see in *Ifnε^-/-^* mice and could be addressed in future studies using *Ifnε^-/-^* conditional knockouts.

Although many cell types respond to IFNε after exogenous treatment with recombinant protein, the cellular targets of IFNε signaling have not been well-defined in models where it is basally expressed. In our scRNAseq analyses of the FRT, we observed ISG expression restricted to cells that also basally expressed IFNε. This ISG expression is dependent on IFNε, as *Ifnε*^−/−^ mice lack ISG expression in the absence of IFNε signaling, suggesting an autocrine role for IFNε. Moreover, in primary human vaginal and uterine epithelial cell lines, we found that IFNε was detectable by ELISA in cell lysates but was undetectable in cell supernatants regardless of viral infection or stimulation. We observed similar results in human enteroids, where IFNε expression could be readily detected by scRNAseq and ELISAs of cellular lysates, but not in supernatants. In contrast to this, we did not see any correlation between *Ifnε* and ISGs in our mouse intestine scRNAseq. Moreover, we observed many differentially expressed genes in the *Ifnε*^−/−^ small intestine, including in cell types that did not express *Ifnε*, indicating that IFNε signaling has effects that extend beyond autocrine signaling. Most notably, *Ifnε*^−/−^ mice lacked entire inflammatory enterocyte populations that expressed *Ifn*λ in WT mice. One explanation for this could be that Ifnε is retained intracellularly and is released from enterocytes as they are shed into the lumen of the intestine as part of normal epithelial turnover and intestinal homeostasis. Future studies will be crucial in elucidating the precise mechanisms by which IFNε initiates signaling and identifying the specific cellular contexts where its autocrine activity plays a protective role, which may provide new insights into its unique function at mucosal barriers.

Unchecked interferon responses can lead to debilitating autoimmune diseases and post infection pathology, highlighting the need for regulation of a constitutively expressed interferon. Retaining IFNε intracellularly within specific epithelial layers vulnerable to infection may prevent aberrant immune activation while still providing basal antiviral protection. However, in cases of cellular damage or viral infection, IFNε could also function as a damage-associated molecular pattern (DAMP). Upon epithelial cell damage or lysis, IFNε may be released into the extracellular space, where it could act in a paracrine manner to signal neighboring mucosal cells or immune cells. This release could enhance local immune responses and alert surrounding tissues to infection or injury, promoting a targeted antiviral response while still limiting widespread immune activation.

Together, our findings underscore the unique role of IFNε in maintaining mucosal immunity as a constitutively expressed IFN within epithelial barriers. By providing baseline antiviral protection while remaining poised for activation upon cellular damage or infection, IFNε serves as both a steady immune defender and a potential DAMP, capable of alerting neighboring cells to pathogenic threats. The enrichment of IFNε in villous-tip enterocytes highlights the importance of spatial positioning within the intestinal epithelium, placing this IFN at an interface where viral entry is most likely to occur. Moreover, the dependence of *Ifnλ3* expression on IFNε reveals a previously unappreciated regulatory axis between constitutive and inducible IFNs that shapes epithelial antiviral programs. This dual functionality helps prevent unwanted immune activation that could lead to autoimmune pathology, while ensuring rapid and localized antiviral responses when needed. Future studies aimed at clarifying the intracellular signaling mechanisms of IFNε and its DAMP-like functions may reveal new therapeutic approaches for enhancing mucosal immunity without triggering systemic inflammation.

## Acknowledgments

This work was supported by R01AI081759 (C.B.C). N.S.H is funded in part by R01AI168107. We thank the Duke University School of Medicine for the use of the Sequencing and Genomic Technologies Shared Resource, which provided RNA-seq services.

## Methods

### Generation of IFNε Knockout Mice Using iGONAD

IFNε knockout (*Ifnε^-/-^*) mice were generated using the Genome Editing via Oviductal Nucleic Acid Delivery (iGONAD) technique, as previously described (Ohtsuka et al., 2018; Skavicus and Heaton, 2023). Briefly, female C57BL/6J mice were mated with C57BL/6J males. On the morning of day E0.7, CRISPR/Cas9-mediated gene editing was performed on pregnant mice by injecting a ribonucleoprotein (RNP) complex consisting of Cas9 protein and two guide RNA (gRNAs) targeting exon 1 (AACGGATTCCCTTCCAATTG[TGG] and AAGTCTTAGCTGCGCTTCAC[CGG], where the PAM sites are indicated with [PAM]) of the *Ifnε* gene into the oviducts. The two gRNAs were designed to flank and induce double-strand breaks, leading to the deletion of a 263 basepair fragment within exon 1, which resulted in a frameshift mutation and knockout of IFNε expression. After injection, the oviducts were electroporated to facilitate the delivery of the ribonucleoprotein (RNP) complex into presumed one-cell stage zygoyes. After gene editing surgery, embryos continued to develop, and pregnant females were allowed to deliver naturally. Genotyping of the resulting pups was carried out by extracting genomic DNA from toe biopsies using a lysis buffer containing 50 mM Tris-base (pH 8.0), 50 mM KCl, 2.5 mM EDTA, 0.45% IGEPAL CA-630 (NP40), 0.45% Tween-20, and 20 mg/mL proteinase K, followed by incubation at 94°C for 10 minutes. PCR amplification was performed with primers flanking the gRNA target sites (forward: TCCCAGAACTGGAGTGGT; reverse: AAGAGCCAACAGGGGATTT) using TITANIUM Taq DNA Polymerase (Takara). The knockout was further confirmed by Sanger sequencing using a primer upstream of the gRNA sites (TCCCAGAACTGGAGTGGT) to verify successful deletion of the *Ifnε* gene. Ifnε^-/-^ mice were backcrossed to C57BL/6J for at least six generations before use in experiments to establish a stable knockout line. Once established, Transnetyx developed a custom qPCR-based genotyping assay to streamline the identification of *Ifnε* knockout alleles in subsequent generations.

### Viruses

HSV-2 (strain 333) was provided by Helen Lazear (University of North Carolina at Chapel Hill) and propagated and titrated using a plaque assay in Vero cells, which were cultured in Dulbecco’s Modified Eagle Medium (DMEM) supplemented with 5% fetal bovine serum (FBS). CVB3 (H3 strain) (Hardy et al., 2004) were propagated and titrated in HeLa cells cultured in Modified Eagle’ s Medium (MEM) supplemented with 5% FBS. For titration, serial dilutions of virus were added to Vero (HSV-2) or CVB3 (HeLa) cell monolayers and incubated for 1 day at room temperature. Following binding, cells were overlaid with a mixture of phenol red-free MEM and 0.5% agarose and incubated for 48-72hrs at 37°C and 5% CO2. The overlay was removed after incubation and plaques were subsequently visualized by staining with crystal violet. Sendai virus (Cantell Strain) was purchased from Charles River (Pl-1, SV).

### HSV-2 infections

8–10-week-old virgin female mice were used for all experiments. For HSV-2 infections females were treated with 2mg of MPA (MedroxyProgesterone Acetate, NDC 66993-270-83) by subcutaneous injection 5 days prior to infection to synchronize females in diestrus. Diestrus was confirmed by vaginal cytology as previously described(Byers et al., 2012) 100 or 10,000 PFU of HSV-2 was given by vaginal inoculation in a 10uL volume. HSV-2 was diluted in sterile PBS. Mice were weighed and scored for pathology daily according to the following criteria: 0 no inflammation; 1 vaginal edema and redness; 2 hair loss or ulceration localized to the vaginal-anal area; 3 severe ulceration, extensive hair loss extending beyond the vaginal-anal area; 4 overt disease signs; 5 paralysis; 6 death. Vaginal washes were taken by pipetting 110uL of PBS in and out of the vagina 5 times. Vaginal lavages were titrated by plaque assay on Vero cells as described above.

### CVB-H3 infections

All CVB-H3 infections were performed in 8–10-week-old male mice. Mice were infected by oral gavage with 1x10^8 pfu in 100uL of PBS. For viral burden experiments mice were sacrificed after 4 days, tissues were harvested and homogenized in DMEM using an Omni International Bead Ruptor Elite (19-042E) at 4m/s for 1 minute using 2.8mm ceramic beads in 2 ml tubes (Omni International 19-628). Viral burdens were quantified using a TCID₅₀ assay on HeLa cells. Serial ten-fold dilutions of virus samples were prepared in complete medium and added to confluent monolayers of HeLa 7B cells in 96-well plates. Cells were incubated for 3 days (37°C, 5% CO₂). Cells were fixed and stained with crystal violet to visualize cytopathic effect (CPE). The dilution at which 50% of wells showed CPE was recorded as the TCID₅₀ endpoint, using the Reed-Muench method to calculate the viral titer.

### Generation of uterine single-cell suspension for scRNASeq

Naturally cycling WT and Ifnε mice in estrus were confirmed by vaginal cytology (Byers et al., 2012). The whole uterus was dissected and placed in 1x DPBS on ice until all tissues were harvested. Uteruses were minced into 1-2mm pieces with dissecting scissors and placed into 2.5mL of D-PBS supplemented with Ca and Mg (Gibco, 14040117). Once all tissues were minced, 2.5 mLs of pre-warmed 2x disassociation media was added. 2x disassociation media was made by dissolving 4mg/mL of Collegnase A (Roche, 10103578001) and 2U/mL of DNAase I (Roche 04536282001) in dispase (Corning 354235). Samples were incubated for 30 minutes at 37 degrees and were shaken vigorously for 30 seconds every 5 minutes. After incubation, enzymes were inactivated by adding 0.5mL of FBS and samples placed on ice. Samples were pushed through 100um filter (VWR 732-2759) using a plunger of a 1mL syringe. Cells were pelleted at 400g for 3 minutes at 4C and resuspended in 1mL of ice-cold PBS. 2ml of RBC lysis buffer (Invitrogen, 00-4333-57) was added for 1 minute with a single inversion to mix, and then 10mLs of 1% FBS-DPBS was added to each sample. Cells were pelleted at 400g for 3 minutes at 4C and resuspended in 500uL of ice cold 1% FBS-PBS. Cells were counted and resuspended at a final volume of 1.5x10^6^ cells/mL in 1% FBS-DPBS. Cells were kept on ice immediately dropped off for scRNAseq processing.

### Generation of intestinal single-cell suspension for scRNASeq

The entire length of the small intestine was dissected from age-matched, 8 week old WT and Ifnε male mice and processed into a single cell suspension as previously described (https://pubmed.ncbi.nlm.nih.gov/36880999/). Briefly, intestinal contents were flushed with 1X HBSS (Sigma H4641), opened longitudinally, and minced into 1mL pieces and then washed twice with 1X HBSS. Tissues were digested for 20 minutes in digestion media: 1X HBSS, 5%FBS (Gibco A56707-01), 5mM EDTA (RPI E14000-250), 1mM DTT (Sigma D0632). Samples were shaken vigorously to help release epithelial cells, and then the supernatant containing epithelial cells was passed through a 100um cell strainer after allowing the residual tissue segments to settle to the bottom of the tube. Samples were spun down at 2000RPM for 5 minutes, resuspended in 1mL of room temperature PBS, and then 2mL of RBC lysis buSer (Invitrogen, 00-4333-57) was added and mixed into the sample by inverting. RBC lysis was performed for 1 minutes at room temperature. 10mL of ice-cold 5%FBS-HBSS was added to each sample, and then samples were strained through a 40uM filter. Cells were counted and resuspended at a final volume of 1.5x10^6^ cells/mL in 5% FBS-HBSS. Cells were kept on ice immediately dropped off for scRNAseq processing.

### Enteroid Growth and Single Cell Preparation

Human fetal enteroid culture and differentiation were based on published protocols for adult stem cells (Beumer et al., 2022; He et al., 2022; Pleguezuelos-Manzano et al., 2020). For expansion, enteroids were passaged by single-cell dissociation in TrypLE (Gibco) at 37°C for 5 min, resuspended in 40 µl Matrigel (Corning) domes, and overlayed with expansion medium (ExM) for 7 days. For differentiation, enteroids were mechanically disrupted in DPBS, resuspended in 40 µl Matrigel domes, grown in patterning medium (PaM) for 16 days and in differentiation medium (DiM) for 3 days. For single-cell analysis, enteroids were expanded (Exp) or differentiated (Dif), scraped into cold DPBS, pelleted 5 min at 400 xg, resuspended in TrypLE, and incubated at 37°C for 15 min. Enteroids were mechanically dissociated by pipetting up and down 30 times, diluted in basal media, pelleted 5 min at 400 xg, resuspended in DPBS with 1% BSA, and filtered through a 30 µm strainer. Basal media contained: 10 mM HEPES, 2 mM L-glutamine, and 100 U/ml penicillin-streptomycin in Advanced DMEM/F12 (Gibco). Expansion media (ExM) additionally contained: 1X B27 (Invitrogen), 1 mM N-acetylcysteine (Sigma), 50 ng/ml mouse EGF (Peprotech), 100 ng/ml mouse Wnt3a, 500 ng/ml mouse R-Spondin, 100 ng/ml human noggin (R&D Systems), 1x N2 (Invitrogen), 10 mM Nicotinamide (Sigma), 100 ng/ml human IGF-1, 50 ng/ml human FGF-2 (Peprotech), 500 nM A83-01 (Tocris), 10 nM PGE2 (R&D Systems), and 10 nM [leu15]-Gastrin I (Millipore). Patterning media (PaM) additionally contained: 1X B27, 1 mM N-acetylcysteine (NAC), 50 ng/ml mouse EGF, 100 ng/ml mouse Wnt3a, 500 ng/ml mouse R-Spondin, 100 ng/ml human noggin, 500 nM A83-01, and 2 ng/ml IL-22 (Peprotech). Differentiation media (DiM) additionally contained 1X B27, 1 mM N-acetylcysteine (NAC), 50 ng/ml mouse EGF, 50 ng/ml BMP-2 and 50 ng/ml BMP-4 (Peprotech).

### Single-Cell RNA-seq Library Preparation and Data Analysis

Publicly available datasets from cycling mouse tissue (Winkler et al., 2024) were downloaded from ArrayExpress (E-MTAB-11491 and from human intestine were downloaded from GEO (accession numbers: GSE185224, GSE171620, GSE125970) (Burclaff et al., 2022; Triana et al., 2021; Wang et al., 2020). Fastq files from all novel single cell RNA sequencing were uploaded to SRA under the bioproject ID PRJNA1329228.

For uterus tissue from WT and *Ifnε^-/-^* mice, single-cell RNA-seq libraries were prepared from ∼10,000 cells/mouse using the 10x Genomics Chromium Single Cell 3’ Reagent Kit (v2 chemistry, Manual Part #CG00052) following the manufacturer’s protocol. Briefly, single-cell suspensions were loaded into the 10x Chromium Controller to capture and barcode individual cells. Libraries were constructed, and sequencing was performed on an Illumina NovaSeq 6000 system (Illumina, San Diego) using an S2 flow cell, providing an average sequencing depth of ∼61,000 reads per cell. After sequencing, raw base call (BCL) files were processed with the 10x CellRanger pipeline (v6.1.2, 10x Genomics) for demultiplexing, alignment to the mouse reference genome (GRCm38), and quantification of gene expression. Quality control and post-processing were carried out using the CellRanger count and aggregate functions to generate gene expression matrices. Data analysis was performed using the Seurat package (Hafemeister and Satija, 2019; Satija et al., 2015) (v4.0) in R. Cells were filtered to include those with at least 700 but no more than 12,000 unique expressed genes. Additionally, cells with over 10% mitochondrial gene content were excluded from further analysis. To correct for batch effects, datasets from individual animals were normalized using the SCTransform() function (v2) in Seurat, and integration was performed using the FindIntegrationAnchors() and IntegrateData() functions. Dimensionality reduction was conducted via principal component analysis (PCA) using the RunPCA() function. Cell clusters were identified using Louvain clustering with the FindClusters() function in Seurat. The optimal clustering resolution was determined by evaluating clustering stability across a range of resolutions (0.2 to 1.0) using the clustree() package, with a resolution of 0.3 selected for downstream analysis. For combined analyses of wild-type (WT) and *Ifnε^-/-^*datasets, previously integrated datasets were merged and further integrated using the Harmony algorithm (Korsunsky et al., 2019) (v1.0) for batch correction and alignment. Differential expression analysis between clusters was performed using the Wilcoxson rank sum test implemented in Seurat’s FindAllMarkers() function. Genes with a log2 fold change > 0.25 and an FDR-adjusted p-value < 0.05 were considered significantly differentially expressed. Marker genes for each cluster were identified by searching for positively enriched genes using the aforementioned criteria.

Human fetal enteroid single-cell RNA-seq libraries were similarly prepared, sequenced and analyzed with minor differences. ∼10,000 cells/sample were targeted, providing an average sequencing depth of ∼61,000 reads per cell. After sequencing, BCL files were processed with the 10x CellRanger pipeline (v7.0.1, 10x Genomics) for demultiplexing, alignment to the human reference genome (GRCh38), and quantification of gene expression. Cells were filtered to include those with at least 3,000 but no more than 10,000 unique expressed genes. Cells with less than 2% or more than 20% mitochondrial gene content were excluded from further analysis. Individual samples were merged and integrated using the FindIntegrationAnchors() and IntegrateData() function into Exp and Dif. Percent mitochondrial genes, feature count, read count, and the difference of G2M and S phase scores were regressed. For combined analysis, Exp and Dif were merged and further integrated using the Harmony algorithm (Korsunsky et al., 2019) (v1.0) for batch correction and alignment.

For intestine tissue from WT and *Ifnε^-/-^* mice, libraries were prepared targeting 20,000 cells/mouse using 10x Genomics v4 GEM-x 3’ chemistry. Libraries were sequenced on a NovaSeq X plus using 2 lanes of a 25B flow cell. Data was analyzed similarly to the uterine tissue, but with some key differences. Data was analyzed using Seurat version 5.3.0 (Hao et al., 2024). Cells were filtered to include those with at least 1,000 but no more than 12,000 unique genes. Cells with greater than 15% mitochondrial gene counts were excluded from further analysis. Individual samples were merged into a split object and normalized and scaled using NormalizeData() and ScaleData() with *Ifne* explicitly retained as a variable feature. WT and Ifn*ε* samples were integrated separately using Harmony Integration and the IntegrateLayers() function. Unique clusters were identified using a resolution of 0.6. To improve detection of lowly expressed transcripts, we applied the ALRA (Adaptively-thresholded Low Rank Approximation) algorithm (Linderman et al., 2022). Differentially expressed genes between WT and *Ifnε^-/-^*clusters were determined using DESeq2.

### Primary cells

Primary human cells were purchased from Lifeline Cell Technologies (FC-0078, FC-0083) and thawed into 24-well or 6-well plates according to manufacturer instructions. At 90% confluency cells were treated with PAMPS or infected at the following concentrations; polyIC float (Invivogen, tlrl-picw) 10ug/mL; polyIC transfection (Inviogen, tlrl-picw) 1ug/mL (0.5ug/well); cGAMP transfection (Invivogen, tlrl-nacga23-02) 10ug/mL; SeV Cantell 20HAU well; HSV-2 333 MOI of 1; LPS float (invivogen, tlrl-eklps) 100ng/mL. cGAMP and polyIC transfected using xTREMEgene9 (Roche, XTG9-RO) according to manufacturer’s instructions.

### qPCR and bulk RNA Sequencing

RNA was extracted from cells using Qiagen Rneasy mini kit (74104) according to manufacter instructions. *IFNε*, *IFNβ*, and *ACTB* RNA levels were detected by qRT-PCR using IDT PrimeTime Std qPCR Assay Primer Probes: *IFNε* Hs.PT.58.4812867; *IFNβ1* Hs.PT.58.39481063.g, ACTB: Hs.PT.39a.22214847 and Itaq one step kit (BIO-RAD 172-5140). qRT-PCR was run using a Biorad Opus. For bulk RNA sequencing, libraries were prepared by the Duke Center for Computational Biology and Genomics (GCB) using a mRNA standed KAPA hyperprep kit. Sequencing was performed an Illumina Novaseq X Plus using 100bp paired end sequencing and targeting 10 billion reads across all samples. Reads were aligned to the human genome using Qiagen CLC Genomics (v20). Differential expression analysis was performed using DESeq2. Briefly, raw count data were imported into R and DESeq2 (Love et al., 2014) was used to normalize counts and identify differentially expressed genes between specified conditions. Genes with an adjusted p-value < 0.05 and log2 ± 2 fold change were considered significantly differentially expressed. Fastq files were uploaded to SRA under the bioproject ID PRJNA1329228.

### ELISA

IFNε and IFNβ protein levels were detected by ELISA (R&D Systems, DY9667-05, DY814-05). IFNε and IFNβ levels were normalized to total protein quantified by Pierce BCA Assay (23227). Cellular lysates were generated using M-PER Mammalian Protein extraction reagent (Thermo Scientific 78501).

### 16S Ribosomal Sequencing

Fecal samples were collected from adult *Ifnε^+/-^ and Ifnε^-/-^* littermates and sent to Microbiome Insights for 16S sequencing and analysis. Specimens were placed into a MoBio PowerMag Soil DNA Isolation Bead Plate. DNA was extracted following MoBio’s instructions on a KingFisher robot. Bacterial 16S rRNA genes were PCR-amplified with dual-barcoded primers targeting the V4 region (515F 5’-GTGCCAGCMGCCGCGGTAA-3’, and 806R 5’- GGACTACHVGGGTWTCTAAT-3’), as previously described (Kozich et al., 2013). Amplicons were sequenced with an Illumina MiSeq using the 300-bp paired-end kit (v.3). Sequences were denoised, taxonomically classified using Silva (v. 138) as the reference database, and clustered into 97%-similarity operational taxonomic units (OTUs) with the mothur software package (v. 1.44.1) (Schloss et al., 2009), following the recommended procedure (https://www.mothur.org/wiki/MiSeq_SOP; accessed Nov 2020). Alpha diversity was estimated with the Shannon index on raw OTU abundance tables after filtering out contaminants. The significance of diversity differences was tested with ANOVA or linear mixed model depending on the study design. To estimate beta diversity across samples, we excluded OTUs occurring with a count of less than 3 in at least 5 % of the samples and then computed Bray-Curtis indices. We visualized beta diversity, emphasizing differences across samples, using Principal Coordinate Analysis (PCoA) ordination. Variation in community structure was assessed with permutational multivariate analyses of variance (PERMANOVA) with treatment group as the main fixed factor and using 999 permutations for significance testing.A dot-dash circle in ordination plot represent for the central of each cluster (Nguyen et al., 2020). All analyses were conducted in the R environment.

---

**Supplemental Figure 1.**
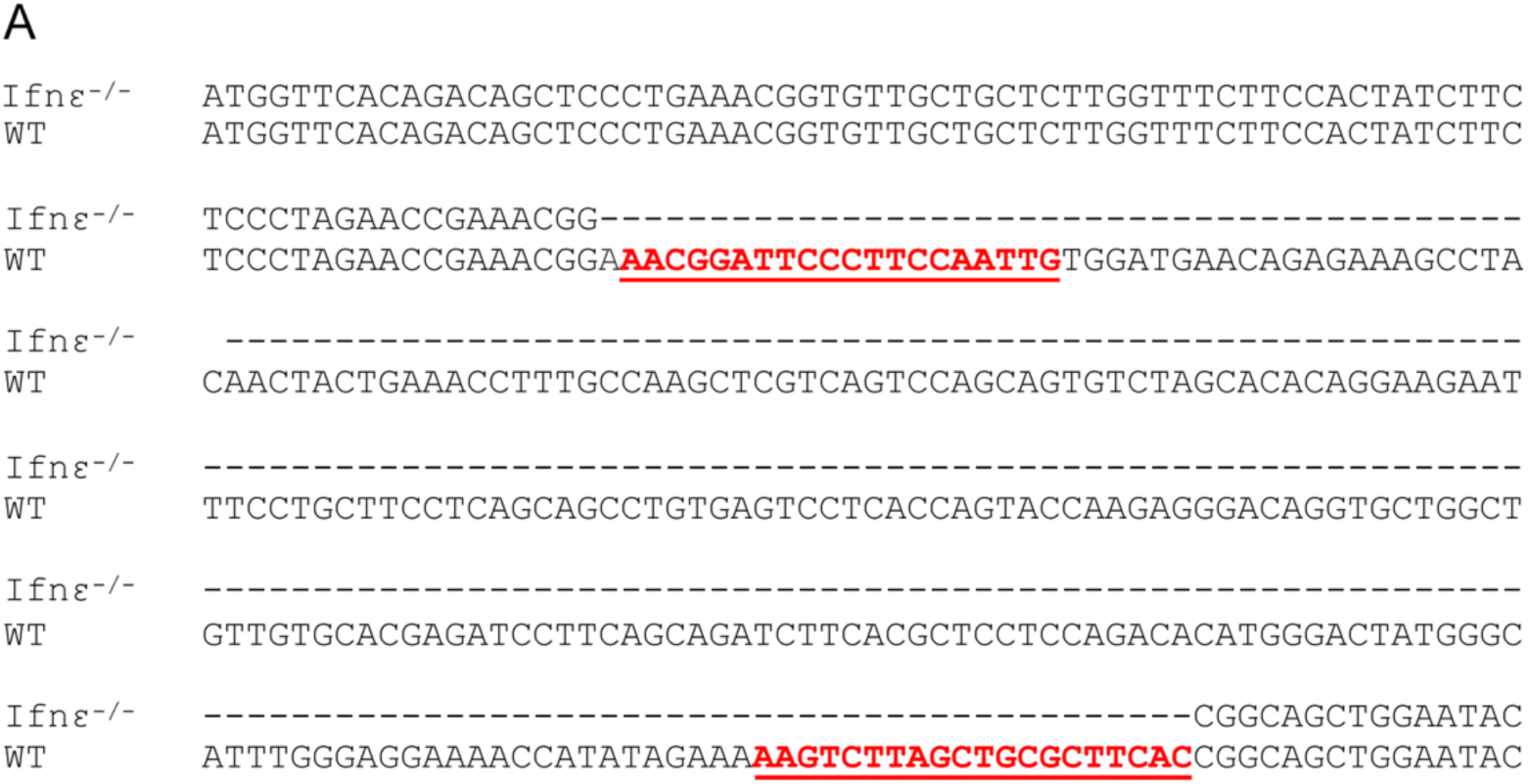
**(A)** Sequence alignment following Sanger sequencing of genomic DNA isolated from an *Ifnε*^-/-^ mouse (top) or wild-type (WT, bottom) mouse. Bold red sequences denote location of gRNAs (gRNA1 and gRNA2). Dashes denote sequence missing in the *Ifnε*^-/-^ mouse.

**Supplemental Figure 2.**
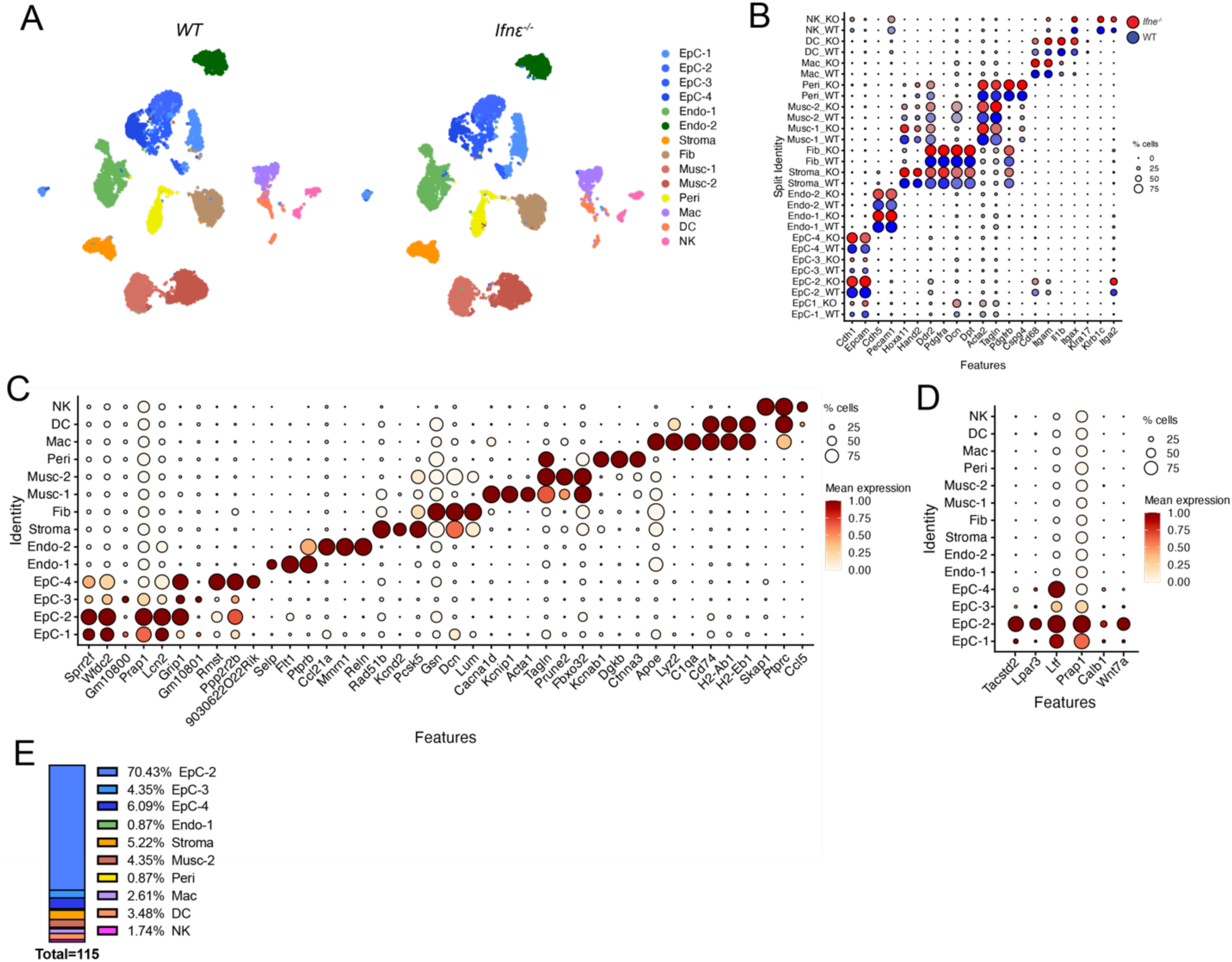
**(A),** UMAP of cell clusters in uterine tissue isolated from WT (left) of Ifnε^-/-^ mice (right) split by genotype. (B), DotPlot of canonical cell markers in WT (blue) or *Ifnε^-^*^/-^ (red) mice. **(C),** DotPlort of cell type markers in the combined dataset from WT and *Ifnε*^-/-^ mice. Scale and key at right. (D), DotPlot of luminal cell markers in WT and *Ifnε*^-/-^ dataset. Key and scale at right. (E), Bar plot indicating the cell clusters associated with differentially upregulated genes (115 total) in WT mice compared to *Ifnε*^-/-^ mice. The percent is shown at right.

**Supplemental Figure 3.**
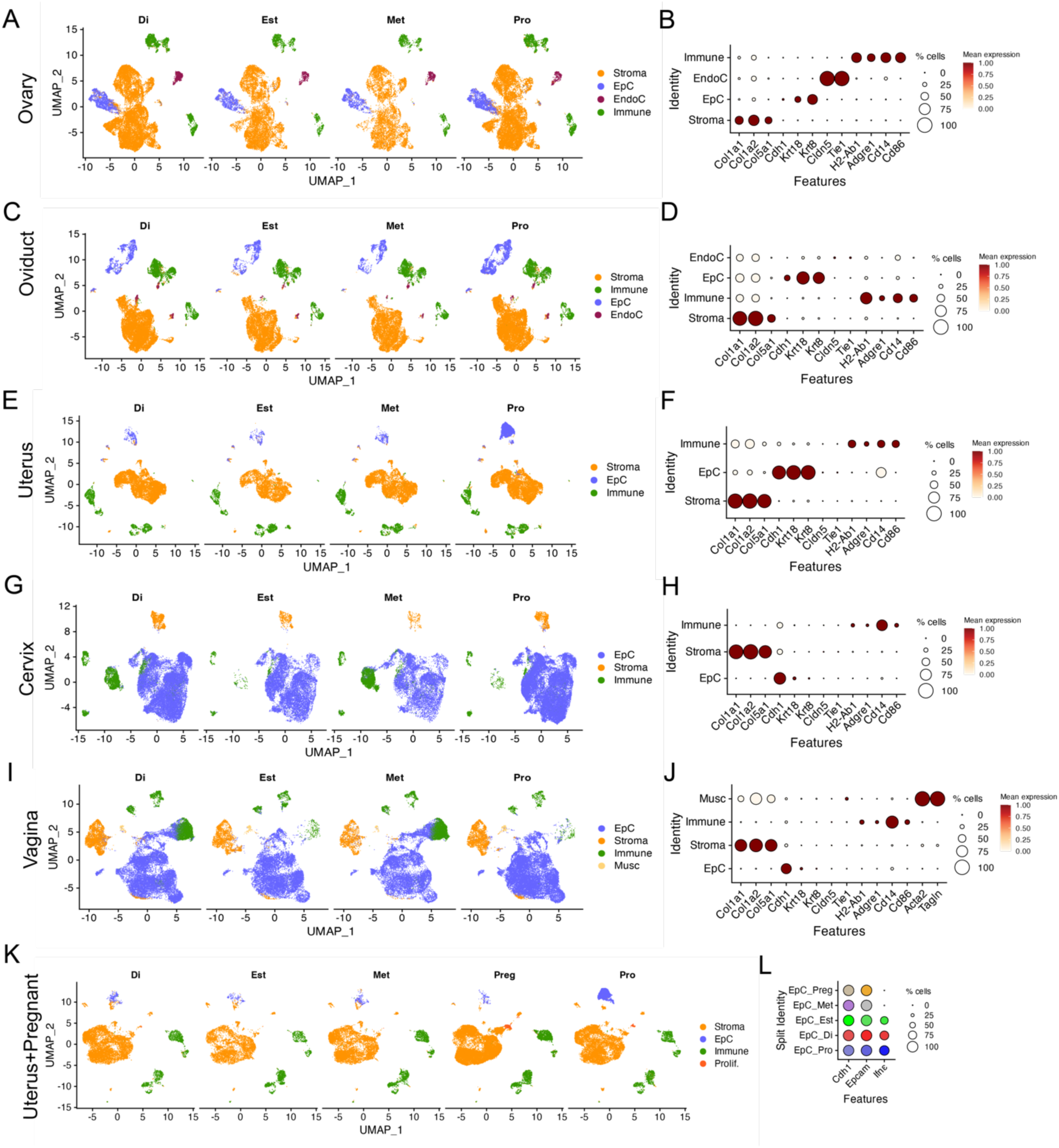
**(A, C, E, G, I)** UMAPS showing clustering of stromal (Stroma, orange), epithelial (EpC, blue), immune (Immune, green), endothelial (EndoC, red), and muscular (Musc, yellow) cells within distinct phases of the estrous cycle the different tissues of the female reproductive tract (FRT) shown at left. **(D, F, H, J),** DotPlot of canonical markers in Oviduct (D), Uterus (F), Cervix (H), Vagina (J), and Uterus (J). **(K),** UMAP of uterine tissue throughout the estrous cycle and pregnant uterus (Preg). **(L),** DotPlot of cadherin-1 (*Cdh1*), *Epcam*, and *Ifnε* in uterine tissue split by phase of the estrous cycle or pregnancy (in orange).

**Supplemental Figure 4.**
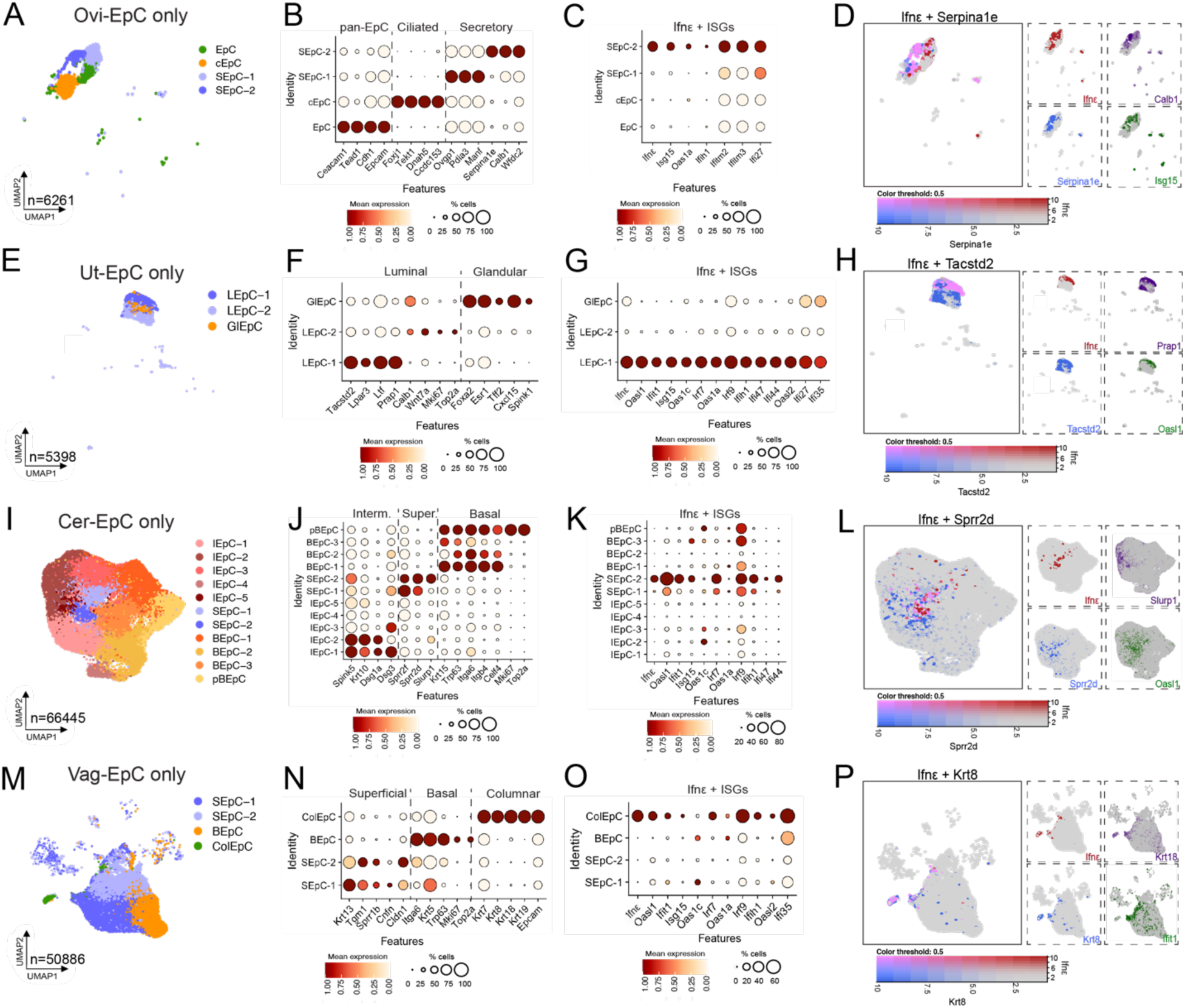
Epithelial cell clusters from Figure 2 were reclustered. **A,E,I,M)** UMAPS showing clustering of epithelial cell types within each tissue. Oviduct **(A)**: general epithelial cells (Epc), ciliated epithelial cells (cEpC), and secretory epithelial cells (SEpC-2). Uterus **(E)**: luminal epithelial cells (LEpC) and glandular epithelial cells (GlEpC). Cervix **(I)**: intermediate epithelial cells (IEpC), superficial epithelial cells (SEpC), basal epithelial cells (BEpC), and parabasal epithelial cells (pBEpC). Vagina **(M)**: superficial epithelial cells (SEpC), basal epithelial cells (BEpC), and Columnar epithelial cells (ColEpC). **B,F,J,N).** Dot plots showing markers of each epithelial subpopulation. **C,G,K,O)** Dot plot showing Ifnε and ISG expression within each cluster. **D,H,L,P)** Feature plots showing expression and coexpression of Ifnε and a top marker for the cluster in which it is expressed.

**Supplemental Figure 5.**
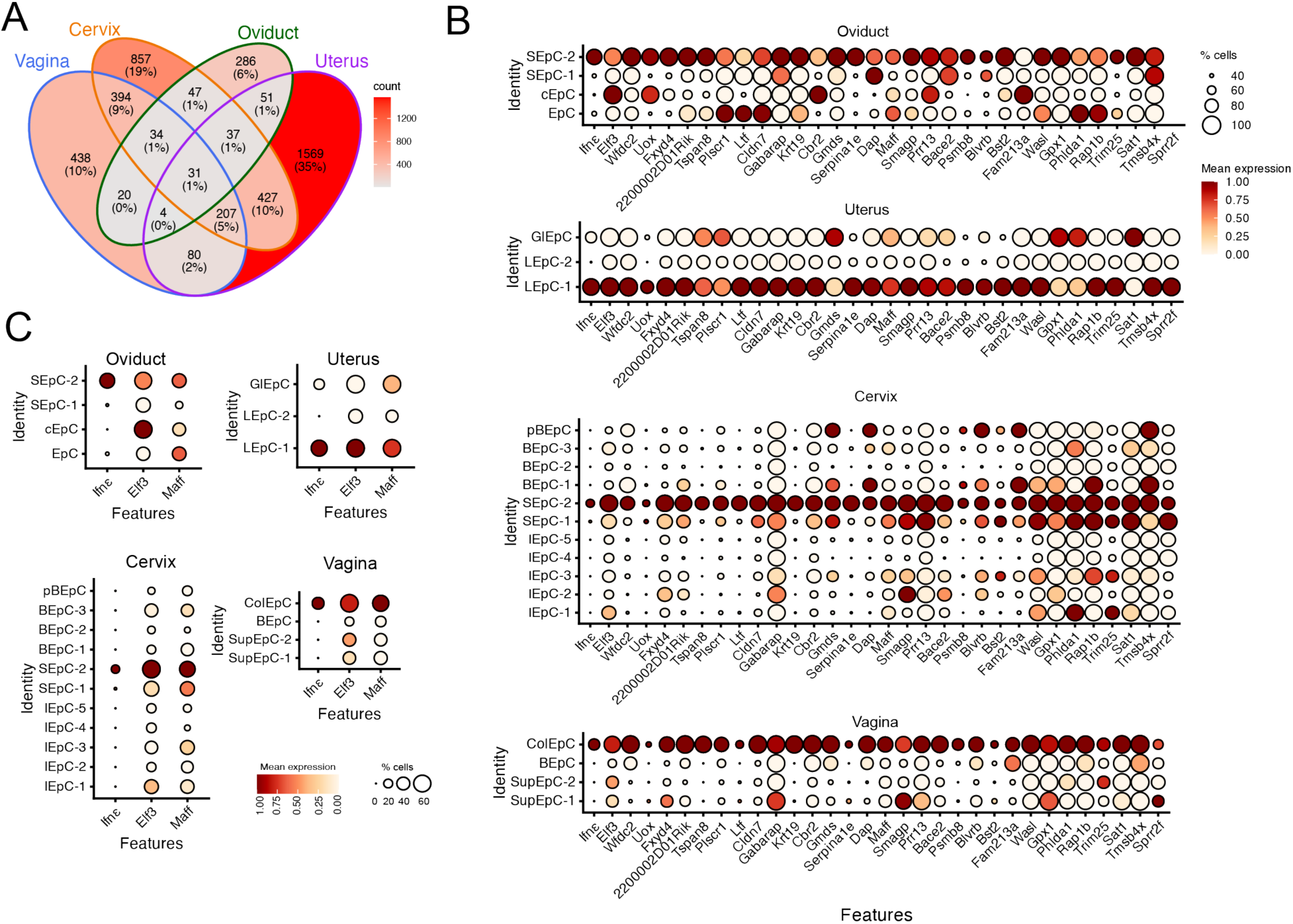
**(A),** Venn diagram denoting the overlap of gene enriched in Ifnε-expressing clusters from Oviduct (green), Uterus (purple), Cervix (orange), and Vagina (blue). Key at right with red indicating the highest number of genes. The percent of overlap is also shown. **(B),** DotPlot of the genes differentially enriched in *Ifnε*-expressing clusters in Oviduct (top), Uterus (second from top), Cervix (second from bottom), and Vagina (bottom). Key and scale at right. **(C),** DotPlot of the expression of *Ifnε*, *Elf3*, and *Maff* in Oviduct, Uterus, Cervix, or Vagina as indicated. Key and scale at bottom.

**Supplemental Figure 6.**
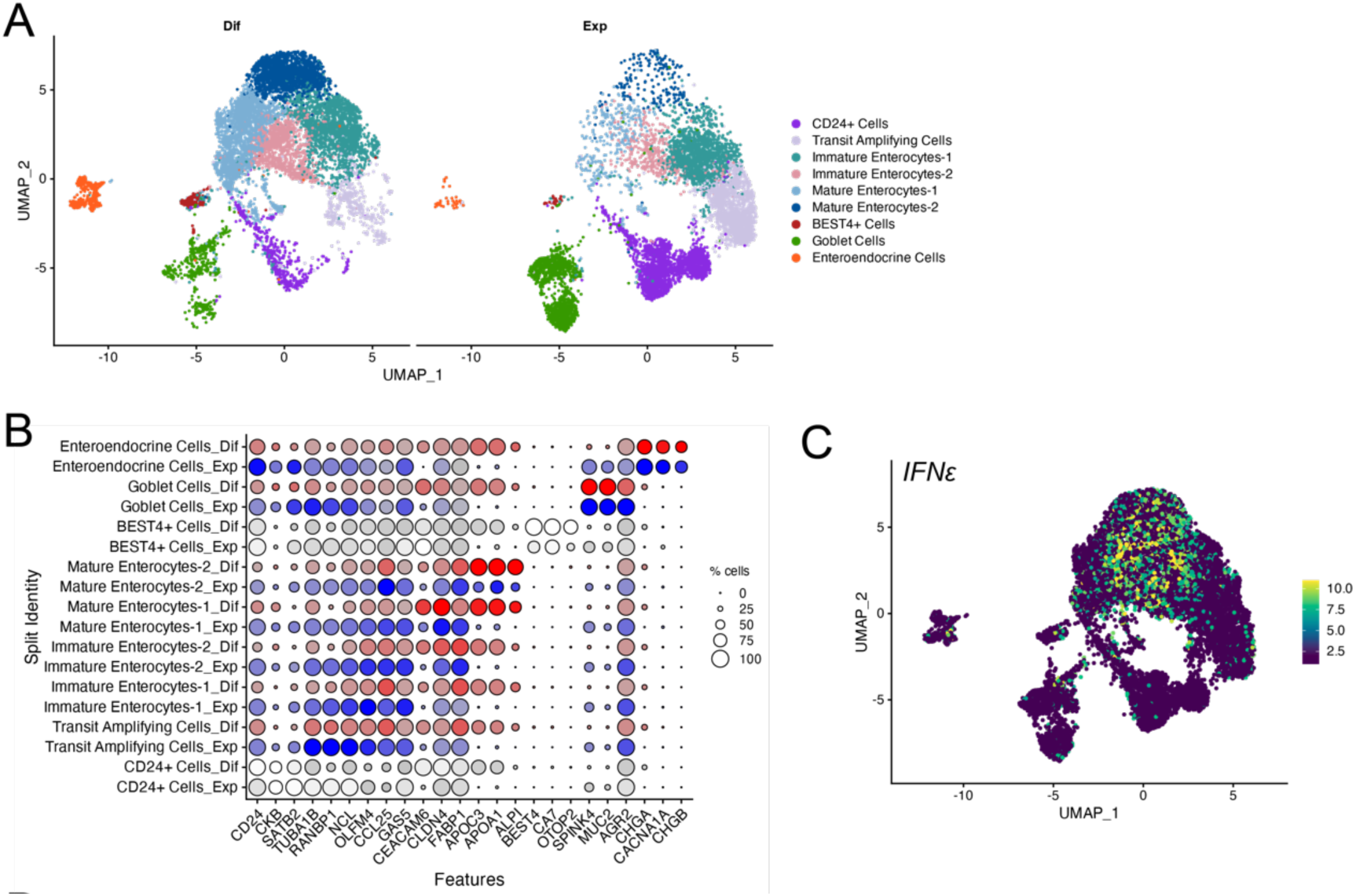
**(A),** UMAP of cell clusters in enteroids cultured under expansion (Exp) or differentiated (Dif) conditions. **(B),** DotPlot of canonical makers in enteroids split by either Dif (red) or Exp (blue) growth conditions. **(C),** FeaturePlot of *IFNE* in enteroids. Scale at right.

**Supplemental Figure 7:**
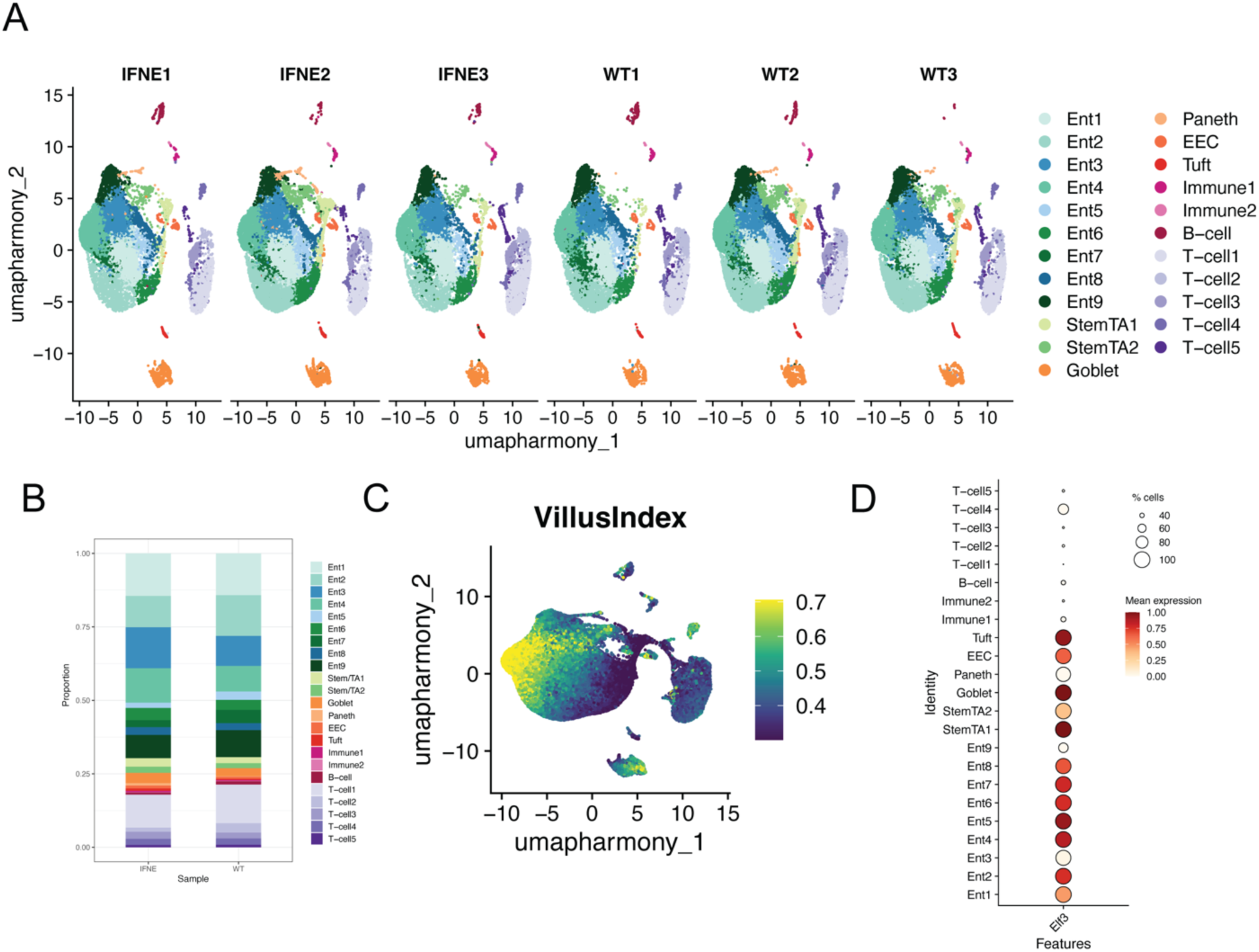
**(A),** UMAP of cell clusters in intestinal tissue isolated from WT or Ifnε^-/-^ mice (right) split by sample. **(B)** Bar plot showing the proportion of cells in each cluster split by WT or Ifnε^-/-^ mice **(C)** Feature plot showing a villus index score for each cell as calculated as follows: (expression top genes/(expression top genes + expression bottom genes). 62 bottom landmark genes and 43 top landmark genes were included that were identified in Moor et al. 2018. **(D)** Dotplot showing proportion and expression levels of *Elf3* in each cluster. Scale and key to the right.

**Supplemental Figure 8:**
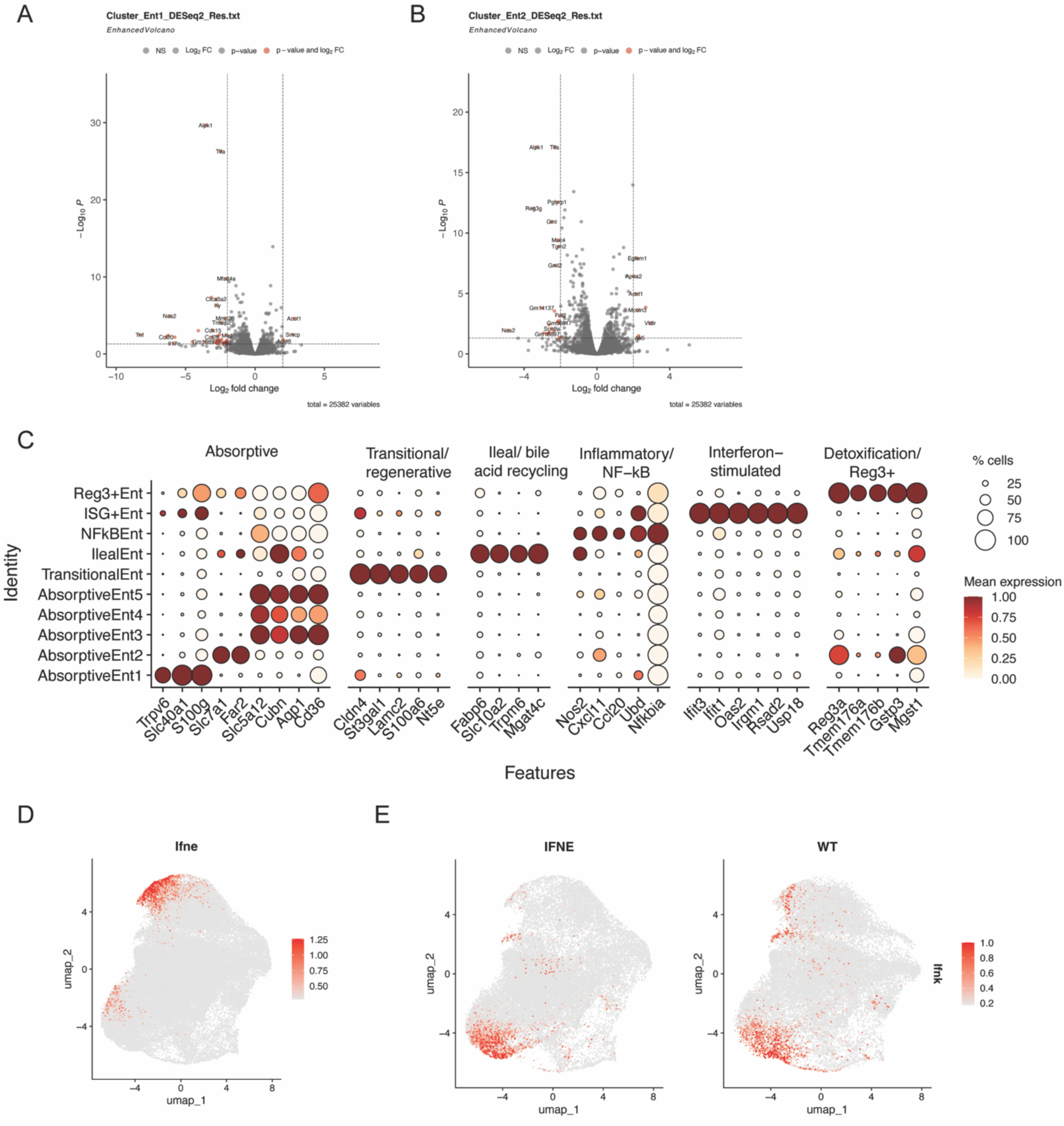
**(A-B)** Volcano plots showing genes differentially expressed in Ifnε^-/-^ samples compared to WT in clusters Ent1 (A) and Ent2 (B). DEGs were calculated by DESeq2. **(C).** DotPlot showing markers used to define types of enterocyte clusters. Key and scale to the right. **(D)** FeaturePlot showing Ifne expression. Scale to the right. **(E)** FeaturePlot showing Ifnk expression split by Ifnε^-/-^ and WT samples. Scale to the right.

**Supplemental Figure 9.**
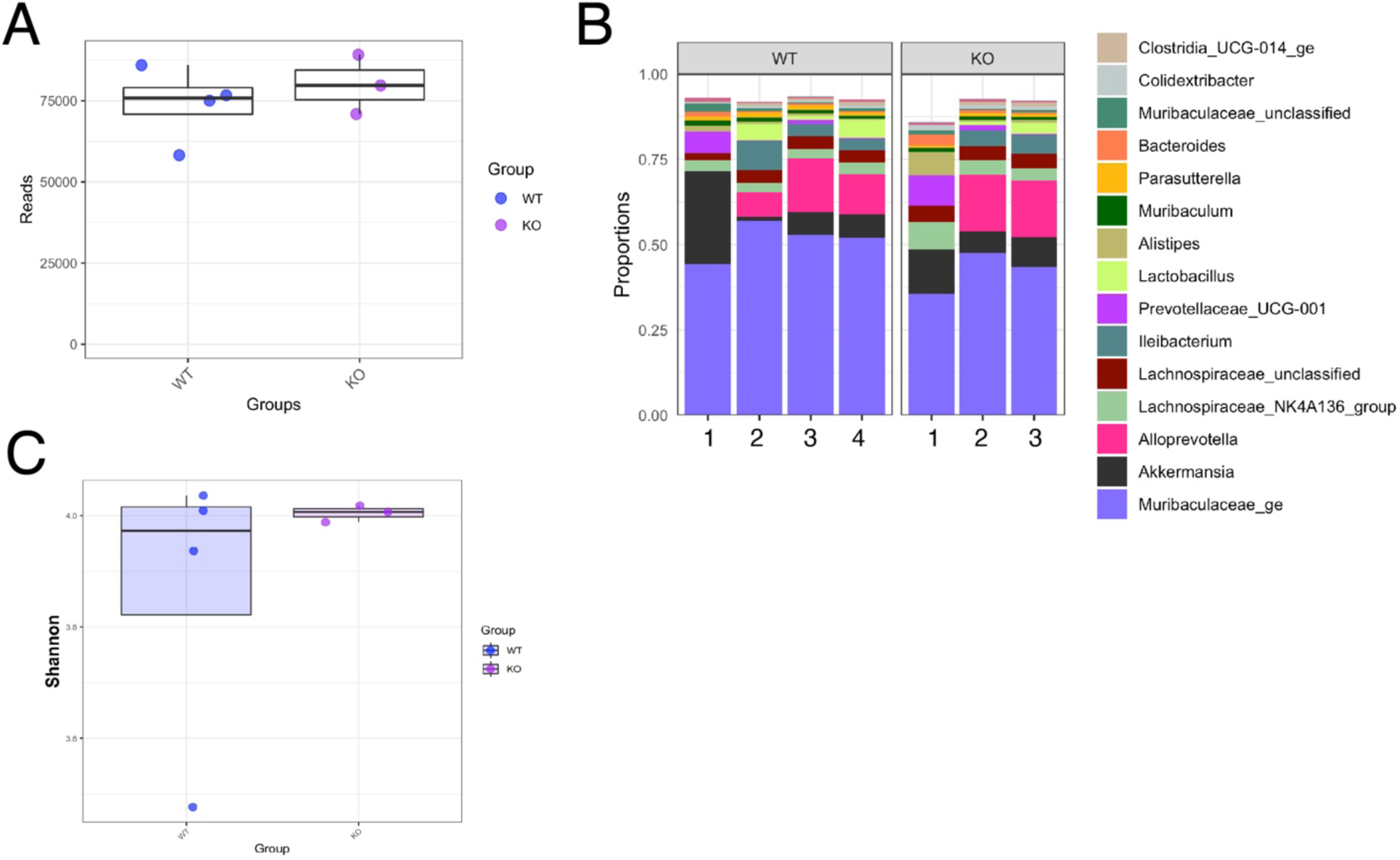
**(A),** Total reads generated from 16S sequencing on fecal samples from *Ifn^+/-^* (WT) and *Ifnε^-/-^* (KO) littermates **(B),** Proportions of OTUs at each taxonomic rank in each sample **(C)** Alpha diversity measured by Shannon Index for *Ifnε^+/-^* (WT) and *Ifnε^-/-^* (KO) littermates.

## Literature Cited

Beumer, J., J. Puschhof, F.Y. Yengej, L. Zhao, A. Martinez-Silgado, M. Blotenburg, H. Begthel, C. Boot, A. van Oudenaarden, Y.G. Chen, and H. Clevers. 2022. BMP gradient along the intestinal villus axis controls zonated enterocyte and goblet cell states. Cell Rep 38:110438.

Bourke, N.M., S.L. Achilles, S.U. Huang, H.E. Cumming, S.S. Lim, I. Papageorgiou, L.J. Gearing, R. Chapman, S. Thakore, N.E. Mangan, S. Mesiano, and P.J. Hertzog. 2022. Spatiotemporal regulation of human IFN-epsilon and innate immunity in the female reproductive tract. JCI Insight 7:

Burclaff, J., R.J. Bliton, K.A. Breau, M.T. Ok, I. Gomez-Martinez, J.S. Ranek, A.P. Bhatt, J.E. Purvis, J.T. Woosley, and S.T. Magness. 2022. A Proximal-to-Distal Survey of Healthy Adult Human Small Intestine and Colon Epithelium by Single-Cell Transcriptomics. Cell Mol Gastroenterol Hepatol 13:1554–1589.

Byers, S.L., M.V. Wiles, S.L. Dunn, and R.A. Taft. 2012. Mouse estrous cycle identification tool and images. PLoS One 7:e35538.

Coldbeck-Shackley, R.C., O. Romeo, S. Rosli, L.J. Gearing, J.A. Gould, S.S. Lim, K.H. Van der Hoek, N.S. Eyre, B. Shue, S.A. Robertson, S.M. Best, M.D. Tate, P.J. Hertzog, and M.R. Beard. 2023. Constitutive expression and distinct properties of IFN-epsilon protect the female reproductive tract from Zika virus infection. PLoS Pathog 19:e1010843.

de Geus, E.D., J.S. Volaric, A.Y. Matthews, N.E. Mangan, J. Chang, J.D. Ooi, N.A. de Weerd, E.M. Giles, and P.J. Hertzog. 2024. Epithelially Restricted Interferon Epsilon Protects Against Colitis. Cell Mol Gastroenterol Hepatol 17:267–278.

Demers, A., G. Kang, F. Ma, W. Lu, Z. Yuan, Y. Li, M. Lewis, E.N. Kraiselburd, L. Montaner, and Q. Li. 2014. The mucosal expression pattern of interferon-epsilon in rhesus macaques. J Leukoc Biol 96:1101–1107.

Deng, S., X. Tian, R. Belshaw, J. Zhou, S. Zhang, Y. Yang, C. Huang, W. Chen, H. Qiu, and S.W. Choo. 2024. An RNA-Seq analysis of coronavirus in the skin of the Pangolin. Sci Rep 14:910.

Fujii, M., M. Matano, K. Toshimitsu, A. Takano, Y. Mikami, S. Nishikori, S. Sugimoto, and T. Sato. 2018. Human Intestinal Organoids Maintain Self-Renewal Capacity and Cellular Diversity in Niche-Inspired Culture Condition. Cell Stem Cell 23:787–793 e786.

Fung, K.Y., E.D. de Geus, L. Ying, H. Cumming, N. Bourke, S.C. Foster, and P.J. Hertzog. 2024. Expression of Interferon Epsilon in Mucosal Epithelium is Regulated by Elf3. Mol Cell Biol 44:334–343.

Fung, K.Y., N.E. Mangan, H. Cumming, J.C. Horvat, J.R. Mayall, S.A. Stifter, N. De Weerd, L.C. Roisman, J. Rossjohn, S.A. Robertson, J.E. Schjenken, B. Parker, C.E. Gargett, H.P. Nguyen, D.J. Carr, P.M. Hansbro, and P.J. Hertzog. 2013. Interferon-epsilon protects the female reproductive tract from viral and bacterial infection. Science 339:1088–1092.

Garcia-Minambres, A., S.G. Eid, N.E. Mangan, C. Pade, S.S. Lim, A.Y. Matthews, N.A. de Weerd, P.J. Hertzog, and J. Mak. 2017. Interferon epsilon promotes HIV restriction at multiple steps of viral replication. Immunol Cell Biol 95:478–483.

Hafemeister, C., and R. Satija. 2019. Normalization and variance stabilization of single-cell RNA-seq data using regularized negative binomial regression. Genome Biol 20:296.

Hao, Y., T. Stuart, M.H. Kowalski, S. Choudhary, P. Hoffman, A. Hartman, A. Srivastava, G. Molla, S. Madad, C. Fernandez-Granda, and R. Satija. 2024. Dictionary learning for integrative, multimodal and scalable single-cell analysis. Nat Biotechnol 42:293–304.

Hardy, M.P., C.M. Owczarek, L.S. Jermiin, M. Ejdeback, and P.J. Hertzog. 2004. Characterization of the type I interferon locus and identification of novel genes. Genomics 84:331–345.

He, G.W., L. Lin, J. DeMartino, X. Zheng, N. Staliarova, T. Dayton, H. Begthel, W.J. van de Wetering, E. Bodewes, J. van Zon, S. Tans, C. Lopez-Iglesias, P.J. Peters, W. Wu, D. Kotlarz, C. Klein, T. Margaritis, F. Holstege, and H. Clevers. 2022. Optimized human intestinal organoid model reveals interleukin-22-dependency of paneth cell formation. Cell Stem Cell 29:1333–1345 e1336.

Jin, S. 2019. Bipotent stem cells support the cyclical regeneration of endometrial epithelium of the murine uterus. Proc Natl Acad Sci U S A 116:6848–6857.

Kellner, M.J., V.M. Monteil, P. Zelger, G. Pei, J. Jiao, M. Onji, K. Nayak, M. Zilbauer, A. Balkema-Buschmann, A. Dorhoi, A. Mirazimi, and J.M. Penninger. 2025. Bat organoids reveal antiviral responses at epithelial surfaces. Nat Immunol 26:934–946.

Korsunsky, I., N. Millard, J. Fan, K. Slowikowski, F. Zhang, K. Wei, Y. Baglaenko, M. Brenner, P.R. Loh, and S. Raychaudhuri. 2019. Fast, sensitive and accurate integration of single-cell data with Harmony. Nat Methods 16:1289–1296.

Kozich, J.J., S.L. Westcott, N.T. Baxter, S.K. Highlander, and P.D. Schloss. 2013. Development of a dual-index sequencing strategy and curation pipeline for analyzing amplicon sequence data on the MiSeq Illumina sequencing platform. Appl Environ Microbiol 79:5112–5120.

Lazear, H.M., J.W. Schoggins, and M.S. Diamond. 2019. Shared and Distinct Functions of Type I and Type III Interferons. Immunity 50:907–923.

Linderman, G.C., J. Zhao, M. Roulis, P. Bielecki, R.A. Flavell, B. Nadler, and Y. Kluger. 2022. Zero-preserving imputation of single-cell RNA-seq data. Nat Commun 13:192.

Love, M.I., W. Huber, and S. Anders. 2014. Moderated estimation of fold change and dispersion for RNA-seq data with DESeq2. Genome Biol 15:550.

Marks, Z.R.C., N. Campbell, N.A. deWeerd, S.S. Lim, L.J. Gearing, N.M. Bourke, and P.J. Hertzog. 2019. Properties and Functions of the Novel Type I Interferon Epsilon. Semin Immunol 43:101328.

Marks, Z.R.C., N.K. Campbell, N.E. Mangan, C.J. Vandenberg, L.J. Gearing, A.Y. Matthews, J.A. Gould, M.D. Tate, G. Wray-McCann, L. Ying, S. Rosli, N. Brockwell, B.S. Parker, S.S. Lim, M. Bilandzic, E.L. Christie, A.N. Stephens, E. de Geus, M.J. Wakefield, G.Y. Ho, O. McNally, S. Australian Ovarian Cancer, I.A. McNeish, D.D.L. Bowtell, N.A. de Weerd, C.L. Scott, N.M. Bourke, and P.J. Hertzog. 2023. Interferon-epsilon is a tumour suppressor and restricts ovarian cancer. Nature 620:1063–1070.

Martinez-Espinoza, I., P.I. Babawale, H. Miletello, N.R. Cheemarla, and A. Guerrero-Plata. 2024. Interferon Epsilon-Mediated Antiviral Activity Against Human Metapneumovirus and Respiratory Syncytial Virus. Vaccines (Basel) 12:

Matsumiya, T., S.M. Prescott, and D.M. Stafforini. 2007. IFN-epsilon mediates TNF-alpha-induced STAT1 phosphorylation and induction of retinoic acid-inducible gene-I in human cervical cancer cells. J Immunol 179:4542–4549.

Mayall, J.R., J.C. Horvat, N.E. Mangan, A. Chevalier, H. McCarthy, D. Hampsey, C. Donovan, A.C. Brown, A.Y. Matthews, N.A. de Weerd, E.D. de Geus, M.R. Starkey, R.Y. Kim, K. Daly, B.J. Goggins, S. Keely, S. Maltby, R. Baldwin, P.S. Foster, M.J. Boyle, P.S. Tanwar, N.D. Huntington, P.J. Hertzog, and P.M. Hansbro. 2024. Interferon-epsilon is a novel regulator of NK cell responses in the uterus. EMBO Mol Med 16:267–293.

McKie, A.T., D. Barrow, G.O. Latunde-Dada, A. Rolfs, G. Sager, E. Mudaly, M. Mudaly, C. Richardson, D. Barlow, A. Bomford, T.J. Peters, K.B. Raja, S. Shirali, M.A. Hediger, F. Farzaneh, and R.J. Simpson. 2001. An iron-regulated ferric reductase associated with the absorption of dietary iron. Science 291:1755–1759.

Miller, D., R. Romero, M. Kacerovsky, I. Musilova, J. Galaz, V. Garcia-Flores, Y. Xu, E. Pusod, C. Demery-Poulos, P. Gutierrez-Contreras, T.N. Liu, E. Jung, K.R. Theis, L.A. Coleman, and N. Gomez-Lopez. 2022. Defining a role for Interferon Epsilon in normal and complicated pregnancies. Heliyon 8:e09952.

Moor, A.E., Y. Harnik, S. Ben-Moshe, E.E. Massasa, M. Rozenberg, R. Eilam, K. Bahar Halpern, and S. Itzkovitz. 2018. Spatial Reconstruction of Single Enterocytes Uncovers Broad Zonation along the Intestinal Villus Axis. Cell 175:1156–1167 e1115.

Nguyen, N.K., E.C. Deehan, Z. Zhang, M. Jin, N. Baskota, M.E. Perez-Munoz, J. Cole, Y.E. Tuncil, B. Seethaler, T. Wang, M. Laville, N.M. Delzenne, S.C. Bischoff, B.R. Hamaker, I. Martinez, D. Knights, J.A. Bakal, C.M. Prado, and J. Walter. 2020. Gut microbiota modulation with long-chain corn bran arabinoxylan in adults with overweight and obesity is linked to an individualized temporal increase in fecal propionate. Microbiome 8:118.

Ohtsuka, M., M. Sato, H. Miura, S. Takabayashi, M. Matsuyama, T. Koyano, N. Arifin, S. Nakamura, K. Wada, and C.B. Gurumurthy. 2018. i-GONAD: a robust method for in situ germline genome engineering using CRISPR nucleases. Genome Biol 19:25.

Pleguezuelos-Manzano, C., J. Puschhof, S. van den Brink, V. Geurts, J. Beumer, and H. Clevers. 2020. Establishment and Culture of Human Intestinal Organoids Derived from Adult Stem Cells. Curr Protoc Immunol 130:e106.

Robinson, C.M., M.A. Woods Acevedo, B.T. McCune, and J.K. Pfeiffer. 2019. Related Enteric Viruses Have Different Requirements for Host Microbiota in Mice. J Virol 93:

Satija, R., J.A. Farrell, D. Gennert, A.F. Schier, and A. Regev. 2015. Spatial reconstruction of single-cell gene expression data. Nat Biotechnol 33:495–502.

Schloss, P.D., S.L. Westcott, T. Ryabin, J.R. Hall, M. Hartmann, E.B. Hollister, R.A. Lesniewski, B.B. Oakley, D.H. Parks, C.J. Robinson, J.W. Sahl, B. Stres, G.G. Thallinger, D.J. Van Horn, and C.F. Weber. 2009. Introducing mothur: open-source, platform-independent, community-supported software for describing and comparing microbial communities. Appl Environ Microbiol 75:7537–7541.

Skavicus, S., and N.S. Heaton. 2023. Approaches for timeline reductions in pathogenesis studies using genetically modified mice. Microbiol Spectr 11:e0252123.

Stifter, S.A., A.Y. Matthews, N.E. Mangan, K.Y. Fung, A. Drew, M.D. Tate, T.P. Soares da Costa, D. Hampsey, J. Mayall, P.M. Hansbro, A. Garcia Minambres, S.G. Eid, J. Mak, J. Scoble, G. Lovrecz, N.A. deWeerd, and P.J. Hertzog. 2018. Defining the distinct, intrinsic properties of the novel type I interferon, IFNɛ. J Biol Chem 293:3168–3179.

Triana, S., M.L. Stanifer, C. Metz-Zumaran, M. Shahraz, M. Mukenhirn, C. Kee, C. Serger, R. Koschny, D. Ordonez-Rueda, M. Paulsen, V. Benes, S. Boulant, and T. Alexandrov. 2021. Single-cell transcriptomics reveals immune response of intestinal cell types to viral infection. Mol Syst Biol 17:e9833.

Van Winkle, J.A., S.T. Peterson, E.A. Kennedy, M.J. Wheadon, H. Ingle, C. Desai, R. Rodgers, D.A. Constant, A.P. Wright, L. Li, M.N. Artyomov, S. Lee, M.T. Baldridge, and T.J. Nice. 2022. Homeostatic interferon-lambda response to bacterial microbiota stimulates preemptive antiviral defense within discrete pockets of intestinal epithelium. Elife 11:

Wang, Y., W. Song, J. Wang, T. Wang, X. Xiong, Z. Qi, W. Fu, X. Yang, and Y.G. Chen. 2020. Single-cell transcriptome analysis reveals differential nutrient absorption functions in human intestine. J Exp Med 217:

Wijayarathna, R., E.D. de Geus, R. Genovese, L.J. Gearing, G. Wray-McCann, R. Sreenivasan, H. Hasan, M. Fijak, P. Stanton, D. Fietz, A. Pilatz, H.C. Schuppe, M.D. Tate, P.J. Hertzog, and M.P. Hedger. 2024. Interferon epsilon is produced in the testis and protects the male reproductive tract against virus infection, inflammation and damage. PLoS Pathog 20:e1012702.

Winkler, I., A. Tolkachov, F. Lammers, P. Lacour, K. Daugelaite, N. Schneider, M.L. Koch, J. Panten, F. Grunschlager, T. Poth, B.M. Avila, A. Schneider, S. Haas, D.T. Odom, and A. Goncalves. 2024. The cycling and aging mouse female reproductive tract at single-cell resolution. Cell 187:981–998 e925.

